# Evolutionary history of inversions in the direction of architecture-driven mutational pressures in crustacean mitochondrial genomes

**DOI:** 10.1101/2020.05.09.085712

**Authors:** Dong Zhang, Hong Zou, Jin Zhang, Gui-Tang Wang, Ivan Jakovlić

## Abstract

Inversions of the origin of replication (ORI) of mitochondrial genomes produce asymmetrical mutational pressures that can cause artefactual clustering in phylogenetic analyses. It is therefore an absolute prerequisite for all molecular evolution studies that use mitochondrial data to account for ORI events in the evolutionary history of their dataset. The number of ORI events in crustaceans remains unknown; several studies reported ORI events in some crustacean lineages on the basis of fully inversed (e.g. negative vs. positive) GC skew patterns, but studies of isolated lineages could have easily overlooked ORI events that produced merely a reduction in the skew magnitude. In this study, we used a comprehensive taxonomic approach to systematically study the evolutionary history of ORI events in crustaceans using all available mitogenomes and combining signals from lineage-specific skew magnitude and direction (+ or -), cumulative skew diagrams, and gene rearrangements. We inferred 24 putative ORI events (14 of which have not been proposed before): 17 with relative confidence, and 7 speculative. Most of these were located at lower taxonomic levels, but there are indications of ORIs that occurred at or above the order-level: Copepoda, Isopoda, and putatively in Branchiopoda and Poecilostomatida+Cyclopoida. Several putative ORI events did not result in fully inversed skews. In many lineages skew plots were not informative for the prediction of replication origin and direction of mutational pressures, but inversions of the mitogenome fragment comprising the ancestral CR (rrnS-CR-trnI) were rather good predictors of skew inversions. We also found that skew plots can be a useful tool to indirectly infer the relative strengths of mutational/purifying pressures in some crustacean lineages: when purifying pressures outweigh mutational, GC skew plots are strongly affected by the strand distribution of genes, and when mutational > purifying, GC skew plots can be even completely (apparently) unaffected by the strand distribution of genes. This observation has very important repercussions for phylogenetic and evolutionary studies, as it implies that not only the relatively rare ORI events, but also much more common gene strand switches and same-strand rearrangements can produce mutational bursts, which in turn affect phylogenetic and evolutionary analyses. We argue that such compositional biases may produce misleading signals not only in phylogenetic but also in other types of evolutionary analyses (dN/dS ratios, codon usage bias, base composition, branch length comparison, etc.), and discuss several such examples. Therefore, all studies aiming to study the evolution of mtDNA sequences should pay close attention to architectural rearrangements.

## Introduction

Mutational pressures predominantly associated with mitochondrial replication often produce strong impacts on the base composition of mitochondrial genomes, which can best be expressed as base composition skews (Reyes et al. 1998a; Xia 2012). In the strand-displacement model of mitochondrial replication, the L-strand is first used as a template to replicate the daughter H-strand, while the parental H-strand is left in a mutagenic single-stranded state for almost two hours (Robberson et al. 1972; Clayton 2000). During this time, spontaneous hydrolytic deamination A and C (into G and T respectively) occurs, generating A-T→G-C and C-G→U-A mutations respectively, the latter of which are much more common (>100 times) (Reyes et al. 1998a; Yasukawa et al. 2006). Over multiple generations, this results in the accumulation of T and G on the H-strand, and A and C on the L-strand. This strand asymmetry, or compositional bias, can be quantified as GC and AT base composition skews, calculated as (G-C)/(G+C) and (A-T)/(A+T) respectively (Perna and Kocher 1995). Due to varying purifying selection constraints, GC skew exhibits greater variability than AT skew, and the greatest differences in GC skew occur at the third codon position and the least skew occurs at the second position (Naylor et al. 1995; Reyes et al. 1998b; Min and Hickey 2007b, a).

Mitochondrial architecture rearrangements that result in an inversion of the replication direction, presumed to be caused by a strand switch of the origin of replication (ORI), change the replication order of the two mitochondrial DNA strands (Hassanin et al. 2005). As this also results in inversed mutational pressures, after a sufficient period of time such events can also result in inversed strand asymmetry (skews) (Hassanin et al. 2005; Wei et al. 2010; Bernt et al. 2013b; Fonseca et al. 2014). This heterogeneity in evolutionary rates can cause artefactual clustering in phylogenetic analyses (Gibson et al. 2005; Hassanin 2006; Min and Hickey 2007a; Rota-Stabelli et al. 2010; Zhang et al. 2019b), and produce misleading signals in analyses of selection pressures acting on protein-coding genes (K_a_/K_s_) (Yang and Nielsen 2000; Zhang et al. 2006). It is therefore an absolute prerequisite for all molecular evolution studies that use mitochondrial data to account for ORI events in the evolutionary history of their dataset.

Cumulative skew plots have been used to indirectly infer ORI events. Most vertebrate mitogenomes have two origins of replication (OR): the strand uncoupling begins at the H-strand OR (O_H_), located within the control region (D-loop or CR), and proceeds unidirectionally until the L-strand OR (O_L_) in the WANCY tRNA cluster is exposed (Sahyoun et al. 2014). As a result, the skew magnitude exhibits a gradient corresponding to the duration of time that the H-strand spends in the mutagenic single-stranded state: the genes closest to the O_L_ in the direction of L-strand replication exhibit the lowest skews, and the genes closest to O_H_ exhibit the highest skews (Reyes et al. 1998a; Faith and Pollock 2003). This produces shift points in cumulative skew diagrams, which have been used to identify the OR (McLean et al. 1998; Touchon et al. 2005; Min and Hickey 2007a; Xia 2012; Sahyoun et al. 2014). However, the actual mitochondrial replication mechanisms are very complex, plastic, often taxon-specific, and only partially understood (Seligmann et al. 2006; Yasukawa et al. 2006; Reyes et al. 2013; Fonseca et al. 2014; Yasukawa and Kang 2018), so skew patterns often produce noisy signals. As a result, identification of the OR is very difficult in mitogenomes with highly rearranged gene order and multiple control regions (Bernt et al. 2013a; Sahyoun et al. 2014). As opposed to vertebrates, mitogenomic architecture in invertebrates is much less conserved and their mitochondrial replication mechanisms are plastic and poorly understood, so CR and replication mechanisms have been identified only in a handful of species (Lavrov et al. 2000; Sahyoun et al. 2014; Gerhold et al. 2014; Lewis et al. 2015; Oliveira et al. 2015).

Over the last few decades, the phylogeny and taxonomy of crustaceans were characterised by a bewildering array of often highly contradictory hypotheses (Schram 2001; Timm and Bracken-Grissom 2015; Lozano-Fernandez et al. 2019), many of which were derived using mitochondrial molecular data, e.g. (Wetzer 2002; Tan et al. 2015, 2019; Timm and Bracken-Grissom 2015; Cheng et al. 2018). A majority of crustaceans exhibit negative GC skews for genes on the mitochondrial ‘majority strand’ (the strand encoding a majority of genes in the ancestral crustacean mitogenomic architecture is named so by convention), but several crustacean lineages, for example most isopods, exhibit inversed (positive) GC skews (Hassanin 2006; Kilpert and Podsiadlowski 2006; Kilpert et al. 2012; Pons et al. 2014; Yu et al. 2018; Zhang et al. 2019b; Zou et al. 2020). As many studies that relied on mitochondrial data to infer the phylogeny of isopods ignored these compositional biases, this produced pervasive topological instability and multiple contradictory phylogenetic hypotheses (Wetzer 2002; Wilson 2009; Hata et al. 2017; Lins et al. 2017; Zou et al. 2018; Zhang et al. 2019b). We have recently shown that in isopods these compositional biases caused by inversions in the strand replication order are so strong that they render mitochondrial data an unsuitable tool for phylogenetics (Zhang et al. 2019b). As these compositional biases can and do affect mitochondrial data-based molecular evolutionary studies in other crustacean lineages as well (Tan et al. 2019), we suspect that a proportion of the contradictory hypotheses for other crustacean lineages may also be attributable to skew-driven phylogenetic artefacts.

Despite this limitation, mitochondrial data, especially complete mitochondrial genomes, can be a suitable tool for inferring lower-level phylogenies and studying evolutionary patterns in crustacean taxa that did not undergo ORI events. While many crustacean lineages exhibit the highly conserved ancestral arthropod mitochondrial architecture (Boore et al. 1998; Cheng et al. 2018; Tan et al. 2018), a number of lineages exhibit exceptionally high plasticity of mitochondrial architecture (Kilpert et al. 2012; Tan et al. 2019; Zou et al. 2020). As this includes common CR duplication events and extremely low levels of conservedness of CR sequences (Pie 2008), no study has ever attempted to systematically infer the evolutionary history of ORI events in crustaceans. It is known that reversal of the strand bias has occurred in crustacean lineages independently multiple times on the basis of fully inversed (e.g. negative vs. positive) skew patterns (Hassanin et al. 2005; Hassanin 2006; Minxiao et al. 2011; Kilpert et al. 2012; Pons et al. 2014; Li et al. 2019; Zhang et al. 2019b), but these data remain scattered throughout the scientific literature, and the exact number of ORI events remains unknown.

Furthermore, we suspected that evolutionary recent ORI events may produce merely a reduction in the skew magnitude, instead of a fully inversed skew; i.e. skew values that have not yet reached equilibrium in their new mutational pressure regime (Rota-Stabelli et al. 2010). Such events could easily have been overlooked by studies of isolated lineages, but they can still generate compositional biases that could produce artefactual results in phylogenetic and evolutionary analyses. A comprehensive taxonomic approach is therefore necessary to systematically study the evolutionary history of ORI events in crustaceans, which in turn is a prerequisite for application of molecular mitogenomic data in evolutionary studies. In this study we studied skews of all available crustacean mitogenomes, and we combined the signals from lineage-specific skew magnitude and direction (+ or -), cumulative skew diagrams, and gene rearrangements to attempt to infer a comprehensive evolutionary history of ORI events in crustaceans.

## Methods

### Dataset and data extraction

We retrieved all 348 crustacean mitogenomes available in the curated RefSeq database (O’Leary et al. 2016); added two Isopoda (Cymothoidae) species with double-inverted skews *Cymothoa indica* (MH396438) and Asotan*a magnifica* (MK790137) (Zhang et al. 2019b; Zou et al. 2020); and a primitive arthropod *Limulus polyphemus* (Chelicerata) with a highly conserved ancestral gene order (Lavrov et al. 2000) as outgroup. To check whether there may be ORIs in taxa not covered by this dataset, we extracted all 991 nominally crustacean mitogenomes from the GenBank, sorted them by GC skew magnitude, selected a cut-off value of GC>-0.2 (this step was conducted to identify taxa with positive and reduced skew magnitudes; as we visually assessed that a vast majority of taxa with conserved ancestral architecture have GC skews <-0.2), compared the taxa with the RefSeq dataset, and selected 16 mitogenomes for additional analyses. Finally, on the basis of ORI events inferred from the above datasets, we retrieved all available mitogenomic data for several lineages with complex skew patterns and multiple ORI events. PhyloSuite (Zhang et al. 2020) was used to retrieve and extract all data, calculate skews, and generate comparative tables and annotation files (taxonomy and gene order) for iTOL (Letunic and Bork 2007). It was also used to update taxonomic data for GenBank files using the default NCBI taxonomy. We set a threshold to 150 bp in PhyloSuite to visualise large non-coding regions (NCR) in mitogenomic gene order maps.

### Phylogenetic analyses

Phylogenetic analyses were conducted to produce guidance trees to help us infer the evolutionary history of ORI events. To make sure that long-branch attraction (LBA) artefacts did not affect our interpretation of the evolutionary history of ORI events, we inspected the overall topology against previous studies of the overall phylogenetic framework for crustaceans (Oakley et al. 2013; Lozano-Fernandez et al. 2019). We used concatenated 13 mitochondrial protein-coding genes (PCGs) to produce a guidance tree of the entire RefSeq dataset (351 species) using Maximum Likelihood (ML) algorithm implemented in IQ-TREE (Trifinopoulos et al. 2016). In our previous study of isopod phylogeny we found that the use of amino acid sequences in combination with the CAT-GTR algorithm, designed to account for compositional heterogeneity and implemented in PhyloBayes-MPI 1.7a (Lartillot et al. 2007), was the most successful strategy for alleviating skew-driven LBA artefacts (Zhang et al. 2019b), so we tested this approach on a reduced dataset (110 species) to assess the prevalence of LBA in our guidance tree. PhyloSuite and its plug-in programs were used to conduct all preparatory steps and ML phylogenetic analysis. Sequences were aligned in batches with ‘--auto’ strategy implemented in MAFFT (Katoh and Toh 2008; Katoh and Standley 2013). trimAI (Capella-Gutiérrez et al. 2009) was used to remove ambiguously aligned regions from the concatenated alignments (using the ‘automated1’ mode). Alignments were concatenating by PhyloSuite, and ML analysis conducted with the automatic model selection (‘-m TESTNEW’ parameter) and 50,000 ‘ultrafast bootstraps’ (Minh et al. 2013). For the PhyloBayes analysis, we used default parameters (burnin = 500, invariable sites automatically removed from the alignment, two MCMC chains). As we failed to achieve conditions considered to indicate a good run, we stopped the analysis when conditions for an acceptable run were met: maxdiff < 0.3 and minimum effective size > 300 (PhyloBayes manual).

### Skew plots, ORI identification and statistical tests

Skews were plotted for the entire mitochondrial majority strand using the algorithm (Xia 2013) implemented DAMBE7 (Xia 2018); where the optimal window and step sizes were automatically inferred according to the GC pattern, unless if specified otherwise. To make plots easier to compare and match to the mitogenomic architecture, average value point of each window was shown at the start of the window, and all GenBank files were rearranged to start with the *rrnS* gene using a custom-made PhyloSuite script. OGDRAW was used to create linear maps of mitogenomic architecture (Greiner et al. 2019). We attempted to infer putative OR locations via shift points in cumulative skew diagrams using the methodology outlined in previous studies (Saito et al. 2005; Breton et al. 2009; Xia 2012). When describing skew plots, we use the terms ‘minimum’ and ‘maximum’ relatively to the presumed direction of mutational pressure; therefore for a species that is evolving under a mutational pressure for a negative GC skew, the maximum will appear as a trough and the minimum as a peak in the skew plot. Similarly, when discussing skew magnitude, we refer to the absolute value (distance from 0). To identify taxa with reduced (but not inversed) skew magnitude, we conducted Welch’s tests using spreadsheets provided in the online version of the Handbook of Biological Statistics (McDonald 2014) and GC skews, as they are best indicators of strand asymmetry (Hassanin et al. 2005; Sun et al. 2018).

## Results

### Tests and calibration of skew plots

To attempt to calibrate the shift points in cumulative skew diagrams (McLean et al. 1998; Touchon et al. 2005; Min and Hickey 2007a; Xia 2012; Sahyoun et al. 2014), and assess whether they can be used to identify the OR in crustaceans, we tested the approach on the human mitogenome, for which the replication mechanism is well-understood. We also tested it on several crustacean/arthropod species that possesses a highly conserved ancestral arthropod gene order (AAGO), including a proto-crustacean ‘living fossil’ *Limulus polyphemus* (Lavrov et al. 2000), and the only arthropod for which the replication mechanism has been described so far (to our knowledge), the fruit fly *Drosophila melanogaster* (Saito et al. 2005). We can relatively safely assume that arthropod mitogenomes with conserved AAGO have reached the mutational equilibrium, as they have been evolving under unidirectional mutational pressures for approximately 400 million years (Lavrov et al. 2000).

In the human mitogenome, O_H_ is located within the control region and O_L_ in the WANCY tRNA cluster, so the genes closest to the O_H_ in the direction of L-strand replication spend the longest time in the mutagenic single-stranded state and exhibit the highest skew magnitude (Faith and Pollock 2003). A previous study proposed that the human mitogenome exhibits two shift points (minima) corresponding to the two origins of replication: a major one corresponding to the location of O_H_ and a minor one corresponding to the O_L_ (Xia 2012). However, that study did not compare the skew plot to the actual mitogenomic architecture. When we did that here, we found that both primary and secondary GC maxima (troughs) were shifted approximately 1500-2000 bp downstream (in the direction of the L strand replication) from the actual OH (Figure 1A). We assume that this can be explained by the large optimal window size of 1948 bp inferred by the DAMBE algorithm, wherein sudden shifts in the skew magnitude are obscured by averaging values over a relatively large segment of the genome. However, as observed before (Xia 2013), small window sizes produce highly noisy plots, from which it is impossible to infer the OR locations (Supplementary File S1: Figures S1-S3). AT skew plot produced a primary maximum that fairly well corresponded to the CR (shifted a few hundred bp downstream), but otherwise it produced a rather noisy plot, with peaks that could easily be misinterpreted as secondary maxima in a mitogenome for which we did not know the exact replication mechanism.

**Figure 1.**
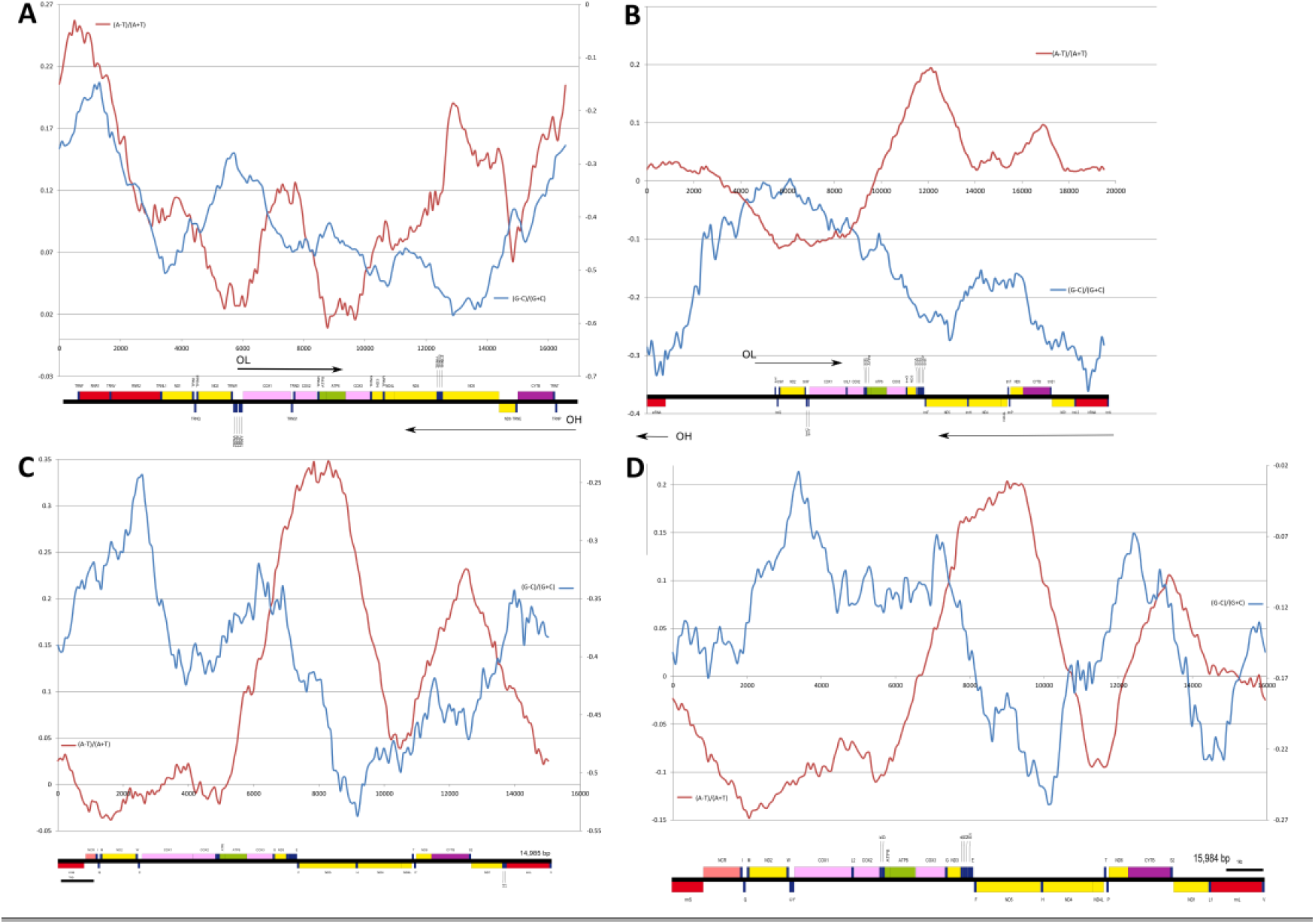
Cumulative skew plots and architecture maps of the mitogenomes of. **(A) *Homo sapiens*** (NC_012920, WS = 1948 bp, SS = 48 bp); **(B) Fruit fly *Drosophila melanogaster*** (NC_024511, WS = 3074, SS = 97), **(C) *Limulus polyphemus*** (WS = 2340, SS = 74); and **(D) *Penaeus monodon*** (Decapoda, WS = 2284, SS = 79). Y-axis shows the skew magnitude and X-axis the mitochondrial sequence (number of bases). When one of the two skews is plotted on the right-hand y-axis, the legend is sown next to the corresponding axis. WS is window size and SS is step size (both in bases). Average values for each window are shown at the starting point. AT skew = (A-T)/(A+T), GC skew = (G-C)/(G+C). The corresponding mitochondrial genome map is shown at the bottom, with the approximate locations of O_L_ and O_H_, and strand replication directions indicated (if known).

Fruit fly possesses a highly conserved AAGO (Boore et al. 1998), wherein the CR is located between the *rrnS* (-strand) and *trnI* (+strand) genes, and both O_H_ and O_L_ are located within that CR. The L-strand (-strand) replication proceeds from O_H_ towards *rrnS* until 97% of the L-strand synthesis is completed, the O_L_ is finally exposed, and the H-strand (major strand) synthesis begins (Saito et al. 2005). Herein, we tentatively treat this as the ancestral replication mechanism for crustaceans as well. GC skew plot for the fruit fly mitogenome fairly well corresponds to the described replication mechanism, but primary maximum was shifted also upstream from the CR (Figure 1B). The primary maximum exhibited two peaks (possibly corresponding to O_H_ and O_L_). A secondary peak, which roughly corresponded to the position of nad4, was mirrored the primary maximum in the AT skew.

Similar to the fruit fly, a segment of the CR adjacent to *rrnS* has been proposed as the origin of replication in a proto-crustacean that also possess a highly conserved AAGO, *L. polyphemus* (Lavrov et al. 2000). However, the GC plot did not produce expected results: the primary maximum roughly corresponded to nad4, and the secondary maximum corresponded to cox2 (Figure 1C). *Penaeus monodon* (Decapoda) (Figure 1D) and *Linguatula serrata* (Ichthyostraca) (Figure 2), two phylogenetically distant crustacean species exhibiting conserved AAGO and negative GC skews, also exhibited the primary GC maxima that approximately corresponded to nad4. However, the secondary maximum for *L. serrata* corresponded to cox2 (due to relatively noisy plot, this is tentative), whereas in *P. monodon* it corresponded to rrnL. This result is in disagreement with the ancestral location of the O_H_ within the CR.

**Figure 2.**
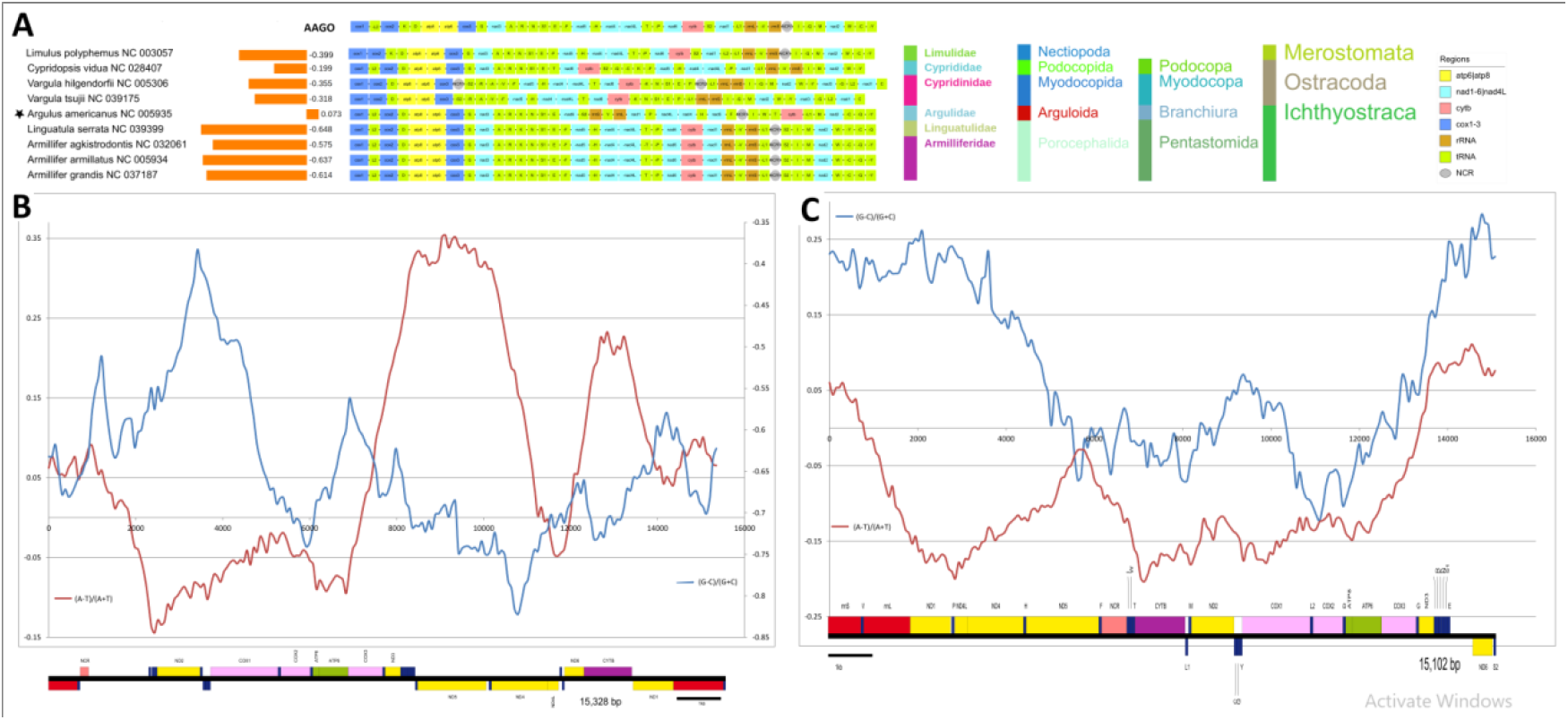
Architecture and skews in the Oligostraca. **(A)** GC skews, gene orders and taxonomy. Star sign indicates an ORI. Ancestral crustacean (arthropod) gene order (AAGO) and *Limulus polyphemus* are also shown; **(B)** Cumulative skew plot and architecture for *Linguatula serrata*. WS = 1437, SS = 76. **(C)** Skew plot and architecture for *Argulus americanus*. WS = 1422, SS = 75. See Figure 1 for other details.

In *L. polyphemus* and *L. serrata*, the AT skew plot produced a primary maximum that roughly corresponded to that of the GC plot (nad5-nad4), and the secondary maximum that approximately corresponded to cytb-nad1. These two maxima closely corresponded to the segments of the genome encoded on the minus strand (note that values averaged over the entire window are shown at the beginning, and not the middle). Indeed, strand distribution of genes fairly well explains the behaviour of the AT skew plot in the fruit fly, and most other crustaceans (see other figures in the manuscript and supplementary data). Intriguingly, the human mitogenome exhibited three AT skew minima, only two of which corresponded to genes encoded on the minus strand, WANCY cluster and nad6-trnE, but the primary minimum corresponded to atp6-cox3 genes, for which we do not have a good explanation.

As regards the GC skews, in the human and fruit fly mitogenomes, despite minor fluctuations corresponding to the locations of genes encoded on the minority strand, skew plots exhibit overall plots that correspond very well to the replication models. However, in the mitogenomes of *L. polyphemus* and *L. serrata*, the primary maxima corresponded to the nad5-nad4L segment encoded on the minority strand, but the other large segment encoded on the minority strand, nad1-rrnS, did not produce such an effect, apart from minor peaks corresponding to nad1 and rrnS respectively. In *Penaeus monodon* (Decapoda), GC skew plot clearly reflected the strand distribution of genes. Due to this inconclusive pattern in the former two species, we further assessed the possibility that origins of replication may be located in the trnF-nad5-trnH-nad4-nad4L segment encoded on the minority strand, or its vicinity. Stem-loops usually serve as origins of replication (Tapper and Clayton 1981), but the mitogenome of *L. polyphemus* has very small intergenic regions, mostly ≤3 bp, except for 9 bp between trnS2 and nad1, which precludes the existence of stem-loops in intergenic regions. We therefore deem it highly unlikely that this segment contains an O_H_, and attribute the primary maxima to asymmetrical purifying mutation pressures produced by genes encoded on a minority strand.

Despite this discordance between shift points in skew charts and the ancestral location of CR, overview of species exhibiting inverted skews indicates that inversions of the mitogenome fragment comprising the ancestral CR (rrnS-CR-trnI) are rather good predictors of skew inversions. Although skew plots appear to produce ambiguous results with respect to the location of the OR, they can produce signals meaningful for our objective of identification of ORI events that did not result in fully inversed skews. Therefore, to infer the evolutionary history of ORI inversions in crustaceans, herein we collated signals from four different criteria: 1) changes in the skew direction and magnitude, 2) strand inversions of the rrnS-CR-trnI region; 3) cumulative skew plots; and in some cases 4) strand inversions comprising nad5-nad4L segment encoded on the minority strand in the AAGO.

## Oligostraca (superclass): Ichthyostraca and Ostracoda (classes)

Oligostraca comprises three classes, but mitogenomes are currently unavailable for the class Mystacocarida.

### Ichthyostraca

The four species available for the subclass Pentastomida all have a highly conserved AAGO and exceptionally high negative GC skews (Figure 2A and B). The only representative of the subclass Branchiura, *Argulus americanus*, has a highly rearranged GO and inverted GC skew (Figure 2C). This was not observed in the original publication of this mitogenome (Lavrov et al. 2004), but Hassanin observed that is has two CRs and elevated evolutionary rate (Hassanin 2006). It exhibits two double-stranded inversions of large segments (i.e., the entire segment changed direction and strand): *F-nad5-H-nad4-nad4L-(T)-P* and *nad1-(L1)-rrnL-V-rrnS*. As almost all of its genes are encoded on the majority strand, both GC and AT skew plots may be informative in this species. Its ancestral CR appears to have been relocated together with trnI between nad5 and cytb. Indeed, there is no NCR between rrnS and trnS2, but there is a 435-bp NCR nearby, between nad6 and trnE, which also almost perfectly corresponds to the GC maximum. AT skew plot produced two maxima: one corresponding to nad1-nad4 and one to the cytb, close to the relocated ancestral CR (trnI-CR). We can conclude only that the mitogenome underwent numerous rearrangements expected to produce an ORI, but we cannot identify the OR with confidence.

### Ostracoda

Among the class Ostracoda, *Cypridopsis vidua* (unpublished) exhibits an rrnS strand switch and a minor decrease in the GC skew magnitude (Figure 2A). However, the conserved architecture of neighbouring segments of mitogenome is indicative of a misannotation artefact. We aligned the *rrnS* of the entire dataset and found that reverse-complement sequence of this gene aligns better with orthologues, which indirectly corroborated our suspicion. It does exhibit a noisy GC skew plot (Supplementary File S1: Figures S4-5), but this is much more likely to be a consequence of a highly rearranged GO than an ORI. Similarly, *Armillifer agkistrodontis* (Ichthyostraca) exhibits strand switches of *rrnL* and *trn*T, both of which could easily be annotation artefacts.

## Branchiopoda, Cephalocarida and Remipedia (classes)

### Branchiopoda

The entire class Branchiopoda exhibits (mostly) highly reduced GC skews (average = -0.0789). Some species have overall GC skews near 0 (−0.005 in *Artemia franciscana*), and two species even have fully inverted (+) GC skews (*Limnadia lenticularis* and *Diaphanostoma dubium*) (Figure 3A). These reduced and inverted skews in the Branchiopoda were either overseen by previous studies (Liu et al. 2017; Luchetti et al. 2019), or merely observed without an in-depth discussion (Bellec et al. 2019). Although this is indicative of inverted architecture-driven mutational pressures, we failed to identify a clear indication of an ORI in the architecture of their mitogenomes, as most of the species have a highly conserved AAGO, with the exception of *Diaphanostoma dubium*. The entire Anostraca clade exhibits a synapomorphic translocation and strand switch (+ to -) of the *trnI* gene. While this suggests that the translocation of this gene may have included the adjacent CR, in disagreement with this observation is the presence of a large NCR adjacent to the *rrnS. Daphnia magna* also exhibits a strand switch for *trnI* gene (+ to -), but without a translocation. *Daphnia carinata* exhibits a GC plot that is indicative of inversed replication order, with positive skews corresponding to the rrnL-rrnS-CR segment (Figure 3B). Somewhat comparable trends were observed in several other species, including the remaining *Daphnia* species, *Triops cancriformis* and *Diaphanostoma dubium*, but *Limnadia lenticularis* (Figure 3C) and *A. franciscana* (Supplementary File S1: Figures S6-S10) exhibited skew plots without a clear trend indicative of replication-driven skews. Taking all this evidence into account, the most parsimonious interpretation is an ORI event that occurred within the CR of the common ancestor of the entire branchiopod lineage, without affecting the neighbouring genes. This produced inverted direction of strand replication compared to the fruit fly (i.e. downstream, as the mitogenome is presented in the Figure 3B), and a mutational pressure towards a positive GC skew. This scenario implies that skews of all species are evolving under unidirectional mutational pressures, but due to the fact that they are not in the equilibrium yet, some GC skews are positive in the fast-evolving segment (around the CR) and negative in the slow-evolving segment. This results in fully inverted (positive) skews in some species, and merely reduced (negative) skews in others. In agreement with this, a previous study found that Anostraca exhibit lower mitochondrial substitution rate than Notostraca and Diplostraca (Luchetti et al. 2019). As Notostraca exhibit the highest negative skews among the branchiopods in our dataset, this indirectly supports our proposed scenario of ancestral negative GC skews that are evolving towards positive skews. As this ORI event would have to be fairly ancient (common ancestor of all branchiopods), incompletely inverted skews imply that evolution of mitogenomes is exceptionally slow in branchiopods.

**Figure 3.**
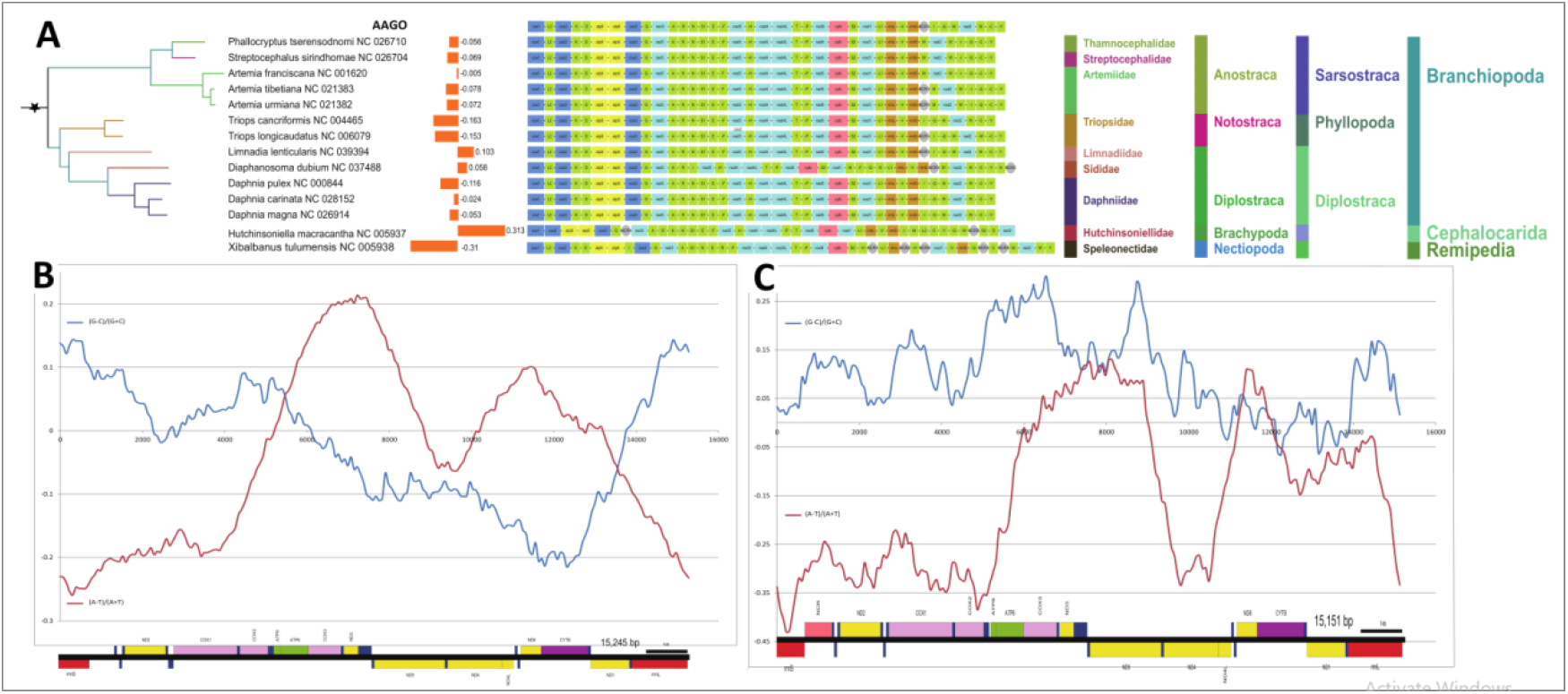
Architecture and skews in the Branchiopoda, Cephalocarida and Remipedia. **(A)** GC skews, gene orders and taxonomy. (**B)** Skew plot for *Daphnia carinata*. WS = 2326, SS = 76. (**C)**. Skew plot for *Limnadia lenticularis*. WS = 885, SS = 76. AAGO is the ancestral gene order, and star sign indicates an ORI. See Figure 1 for other details.

### Cephalocarida

Inverted skew in the only available Cephalocarida species, *Hutchinsoniella macracantha*, was observed before (Hassanin 2006), but not its exceptional magnitude: the overall GC skew of 0.313 is the second highest positive value in the entire dataset after *Polyascus gregaria* (Thecostraca) (Supplementary File S2). It exhibits a large number of rearrangements (Figure 3A), two large NCRs in noncanonical positions, and a strand switch of *trnI* (+ to -). Intriguingly, there is no intergenic region between rrnS and trnI, but there is a 118 bp NCR between trnV and rrnS, and there are two additional large NCRs in the mitogenome. Therefore, we can conclude that there some of the multiple architecture rearrangements and strand switch events resulted in a translocation and inversion of the CR, which produced an ORI.

### Remipedia

Represented by only one species, *Xibalbanus tulumensis* (syn. *Speleonectes tulumensis*), with a rearranged GO, but the skew is standard, and none of the rearrangements include strand switches of genes putatively associated with the ancestral CR (Figure 3A).

## Thecostraca (class)

The entire class exhibited a strongly reduced average skew magnitude compared to decapods with conserved AAGO (−0.18 vs. -0.27 respectively), but all species apart from *Chthamalus* sp. (Chthamalidae) and *Polyascus gregaria* had negative GC skews and the ancestral arrangement of rrnL-V-rrnS-CR, so there is no clear indication of an ancestral ORI (Figure 4A). Both available *Chthamalus* species (*antennatus* and *challenegeri*) exhibited a double-stranded inversion (both strand and direction change) of a large genomic fragment comprising 17 genes and the ancestral CR, *nad5-H*-*nad4-nad4L-P-T-nad6-cytb-S2-Y-C-nad1-L1-rrnL-V-rrnS-CR-K* (Chen et al. 2019), which explains the inverted GC skew (0.065 and 0.092). As the other available Chthamalidae genus, *Notochthamalus* (*scabrosus*), did not exhibit this inversion, it appears that the ORI occurred in the ancestor of the genus *Chthamalus*. This skew inversion was not observed before (although (Chen et al. 2019) calculated skew values, they did not observe that they are inverted in comparison to most other Thecostraca).

**Figure 4.**
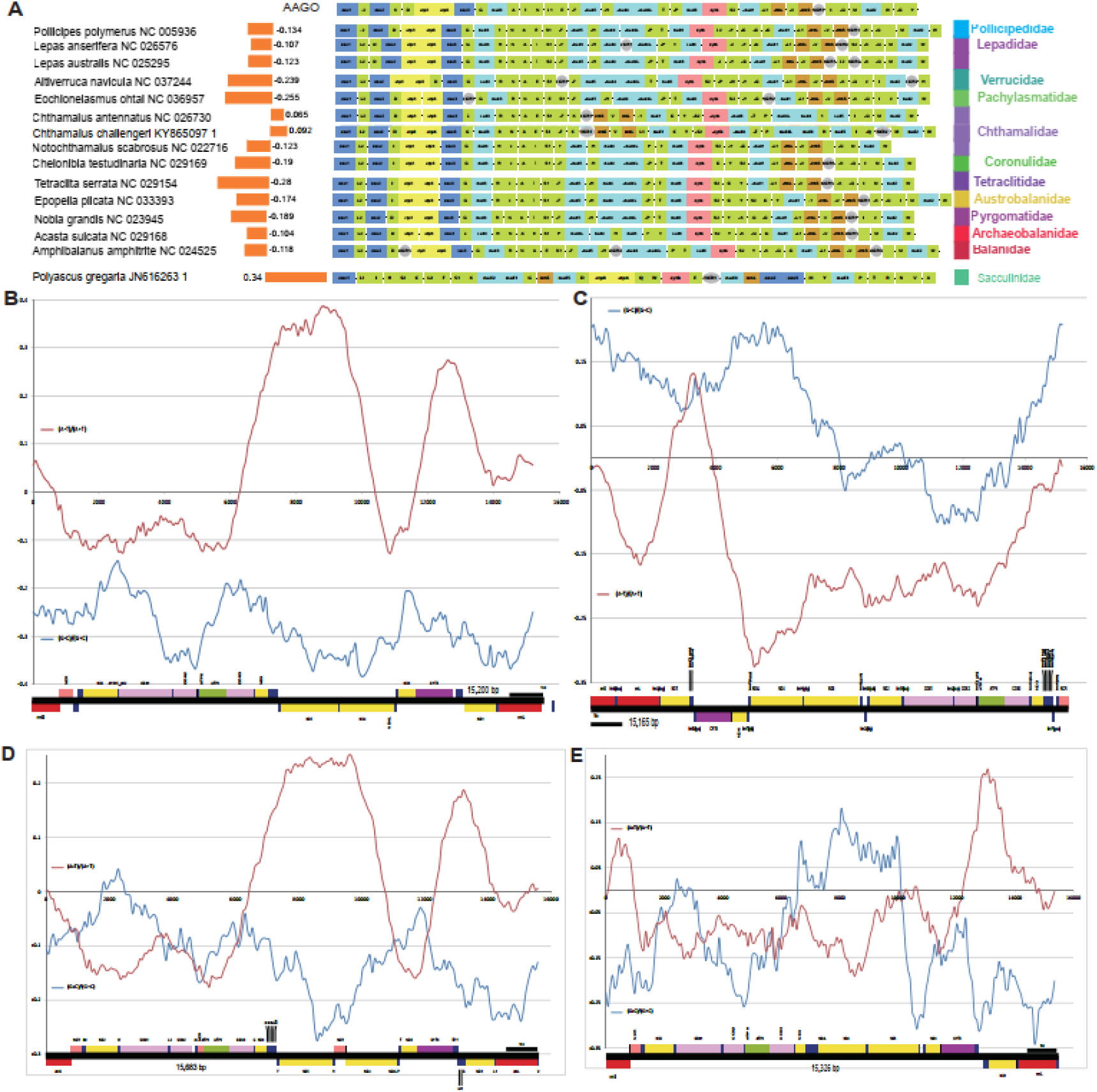
Architecture and skews in the Thecostraca. **(A)** GC skews, gene orders and taxonomy. AAGO is the ancestral gene order. Cumulative skew plots for: (**B)** *Tetraclita serrata*. WS = 1426, SS = 76. **(C)** *Chthamalus antennatus*. WS = 1752, SS = 76. **(D)** *Amphibalanus amphitrite*. WS = 1500; SS = 76. Global GC Skew = -0.11848, Global AT Skew = 0.00565. **(E)** *Acasta sulcata*. WS = 1500 (manually set); SS = 76. Global GC Skew = -0.10381, Global AT Skew = -0.04262. See Figure 1 for other details.

### Rhizocephala

The mitogenome of the only representative of the superorder (or infraclass, depending on the source) Rhizocephala, *Polyascus gregarius* (syn. *gregaria*), exhibits a unique architecture among the crustaceans, with all genes encoded on the plus strand (Yan et al. 2012). This explains not only the ORI, but also the highest positive GC skew among all crustaceans: 0.34 (Supplementary File S2). Its inverted skew was not observed in the original publication. Unfortunately, this is the only representative of the entire superorder, so we cannot make any further guesses about the phylogenetic placement of this unique arrangement and the associated ORI. We urge sequencing of further mitogenomes belonging to this lineage, as they may represent a uniquely interesting model for studying the evolution of mitochondrial architecture.

### Thoracica

As regards remaining species, all of which belong to the superorder (or infraclass) Thoracica (barnacles), we propose at least two tRNA gene rearrangements as synapomorphic for the entire clade: the ancestral *trnT*(+)-*trnP*(−) rearrangement into *trnP*(−)-*trnT*(+), and translocation of trnI (adjacent to *rrnS* in the AAGO).

Rearrangement of trnI and decreased skew magnitude suggest a possibility that a segment of the CR containing the O_L_ adjacent to it may have been translocated as well, but we identified a large NCR in the ancestral crustacean location in most taxa, and there are no notable intergenic regions adjacent to trnI in families that exhibit the putatively ancestral Thecostraca arrangement (Q-I-M), Tetraclitidae, Austrobalanidae and Coronulidae. A species with a (putatively) standard ancestral GO for Thecostraca, *Tetraclita serrata*, exhibits a GC skew plot with a very weak trendline that cannot be interpreted unambiguously (Figure 4B), but it indicates very weak mutational pressures. The skew plot for *Amphibalanus amphitrite* (Figure 4D), exhibits an almost identical profile, but the value corresponding to the putative CR dips into positive values, which indicates that these species may indeed have undergone an ORI, but their mitogenomes are evolving very slowly. However, that scenario is in apparent disagreement with the positive GC skew, a similar skew profile, and rearrangement indicative of ORI in *Chthamalus antennatus*. This skew plot exhibits a skew maximum corresponding to the CR, followed by a steady downward trend, so the segment of the graph that corresponds to cox1-cox2 dips into negative values, and finally steep increase back to the maximum between atp6 and CR (Figure 4C). Negative values correspond to the segment farthest from the OH in this scenario, and not to a change in the trendline, so they can be interpreted as incomplete skew inversion. Three (out of five) species within the family Balanidae (*Acasta* and *Megalbanus* genera) exhibited a double-stranded inversion (changed strand and direction) of a large fragment: *F-nad5-H-nad4-nad4L-P* (−to +). This merely produced slightly reduced overall skews compared to other related species. GC skew plot for *Acasta sulcata* indicates either that the rearrangement was recent, or that the mitogenomes are evolving extremely slowly, as the inverted segment still exhibits positive skew values (Figure 4E). To explain in more detail, in the ancestral arrangement, plus strand had negative GC skews, and minus strand had positive GC skews. As the two strands became inverted, now positive GC skews appear on the plus strand. Therefore, we can infer with confidence only that mitogenomes of most of the Thoracica species possess very low negative GC skews, and evolve under very weak mutational pressures. We cannot infer with confidence the direction of these pressures; i.e. we cannot infer whether the common ancestor of this lineage may have undergone an ORI, or whether the low skew magnitude is a product of exceptionally low mutational pressures.

## Copepoda (class)

This entire class exhibits destabilised mitogenomic architecture, with the orders Poecilostomatida and Cyclopoida exhibiting negative skews, Siphonostomatoida positive, whereas Harpacticoida and Calanoida exhibited mixed positive and negative skews (Figure 5A). Calanoida also exhibited some of the most highly disrupted mitogenomic architectures in the entire crustacean dataset. Due to their multiple large NCRs, with reported sizes >20 Kbp (Weydmann et al. 2017), *Calanus* spp. mitogenomes might be the largest among all crustaceans, along with the decapod genus *Metanephrops spp*. (≈20.6 Kbp; Supplementary File S2). Inverted skews in isolated copepod taxa have been observed before (Hassanin 2006; Minxiao et al. 2011), but due to their highly disrupted architecture, it is difficult to identify the rearrangement events that produced ORIs. As Poecilostomatida and Cyclopoida (negative skews) are closely related sister-orders highly derived within the Copepoda (Khodami et al. 2017), tentatively we accept the hypothesis of (Minxiao et al. 2011) of an ORI in the common ancestor of the entire clade. This appears to have been followed by a subsequent ORI in the common ancestor of Poecilostomatida+Cyclopoida, resulting in D-I skews. The genus *Tigriopus* (Harpacticoida) exhibited an intriguing skew pattern: two species had some of the highest positive GC skews (≈0.27) in the entire dataset (Supplementary File S2), whereas *T. kingsejongensis* had a high negative skew of -0.16. The latter species exhibits a highly rearranged GO compared to the former two species, including the strand inversion of rrnS (minus to plus strand). Assuming an ancestral ORI (we do not reject alternative hypotheses), this implies an ORI and D-I skew in *T. kingsejongensis*. This ORI was not observed previously. As such a large difference in skews among congenerics is puzzling, we compared their cox1 barcodes, and found only 75% similarity. This suggests that they may have split from the common ancestor a long time ago and/or that they are evolving under elevated mutational rates. In partial agreement with this, copepods have some of the highest mitochondrial mutation rates among all bilaterian animals (Bernt et al. 2013a).

**Figure 5.**
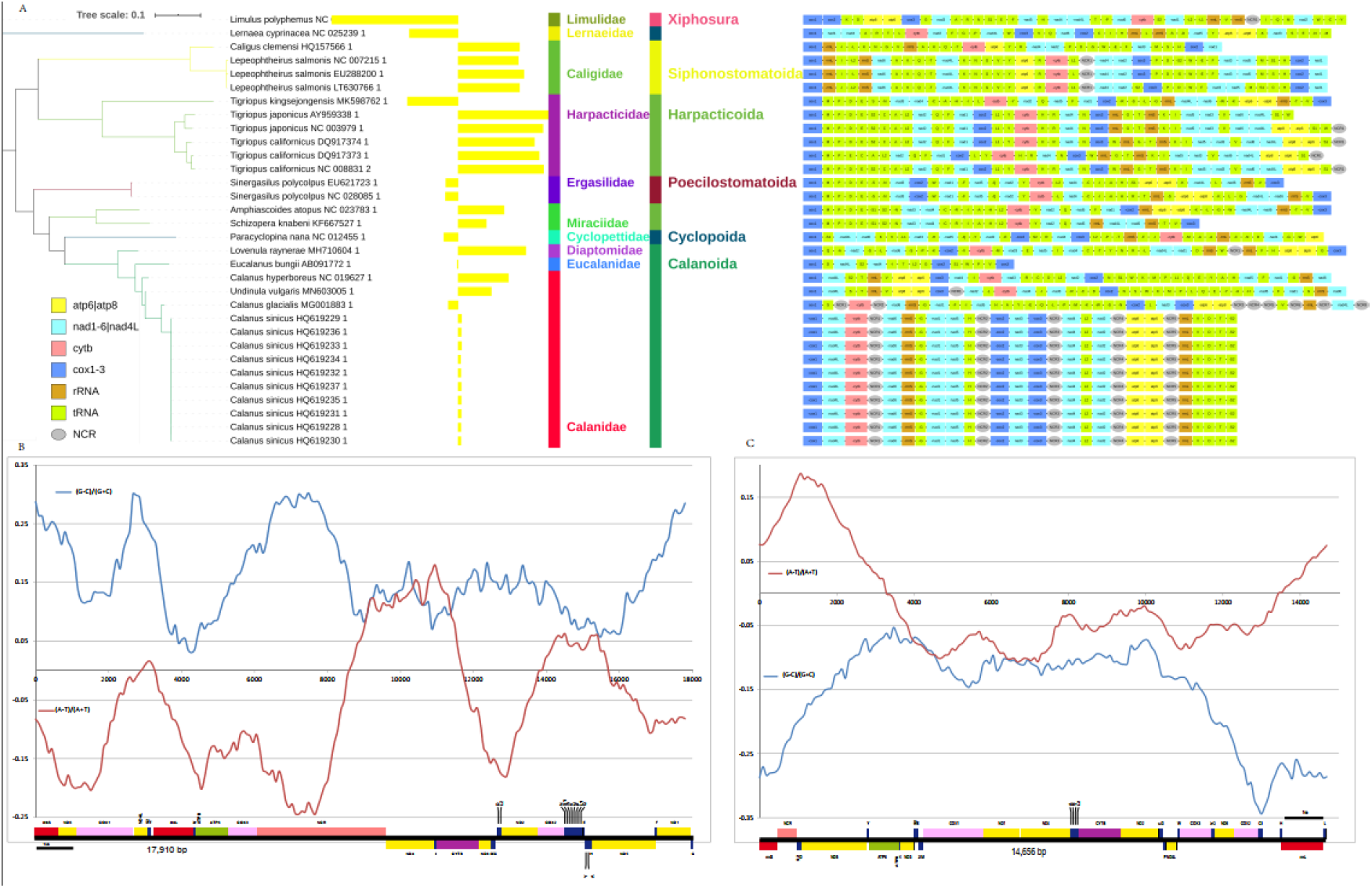
Architecture and skews in the Copepoda. **(A)** GC skews, gene orders and taxonomy. (**B)**. Skew plot for *Calanus hyperboreus*. WS = 1500 (manually set), SS = 89. (**C)**. Skew plot for *Lernaea cyprinacea*. WS = 2973, SS = 89. AAGO is the ancestral gene order, and star sign indicates an ORI. See Figure 1 for other details.

Weydmann et al. found that *Calanus* spp. mitogenomes exhibit uneven skew patterns, switching between positive and negative values, and suggested independent replication from different NCRs and/or frequent change of the mutational pressure associated with mitochondrial DNA replication (Weydmann et al. 2017). We suspect that a large part of the explanation for this lies in the observation discussed in the previous section: frequent strand switches of genes produce noisy skew plots. Intriguingly, the algorithm implemented in DAMBE inferred very small optimal window sizes for the mitogenomes of Copepoda (Supplementary File S1: Figures S11-18). An exception is *Lernaea cyprinacea* (Cyclopoida), which exhibits a negative GC skew, the ancestral arrangement of putative CR adjacent to the rrnS(−), and a GC skew maximum that roughly corresponds to that segment of the mitogenome (spanning cox2-trnC1-NCR-H-rrnL-L-rrnS-NCR) (Fig 5C). We urge sequencing of further copepod (particularly Calanoida) mitochondrial genomes, as they seem to be exceptionally interesting from the aspect of mitogenomic architecture evolution.

## Isopoda (class Malacostraca: subclass Eumalacostraca: superorder Peracarida)

Inversed skew as autapomorphic trait for all isopods apart from the putatively basal Asellota has been proposed almost a decade ago (Kilpert et al. 2012), but recently we showed that Cymothoidae and Corallanidae underwent an additional ORI and exhibit double-inverted (D-I) skews (Zhang et al. 2019b). Isopods generally have highly destabilized, hypervariable GOs, with almost all species sequenced so far exhibiting a unique GO (Zou et al. 2020), and with a large number of genes exhibiting strand switches compared to the AAGO (Figure 6A). Notably, all available isopod mitogenomes are characterised by the minus to plus strand switch of *rrnS*, and *rnI* is missing from many mitogenomes. We can therefore infer with some confidence that this ancestral isopod strand inversion included the CR, and produced an ORI. However, as Asellota also exhibit this rearrangement, this invalidates the hypothesis that the ancestral ORI took place after the (basal) Asellota branched off from the isopod lineage (autapomorphy) (Kilpert et al. 2012; Zhang et al. 2019b), and indicates that ancestral ORI took place in the common ancestor of all isopods (synapomorphy). This scenario requires two subsequent independent ORI events in isopods: one in the common ancestor of Asellota, and one in the common ancestor of Cymothoidae and Corallanidae, both resulting in D-I skews (Figure 7).

**Figure 6.**
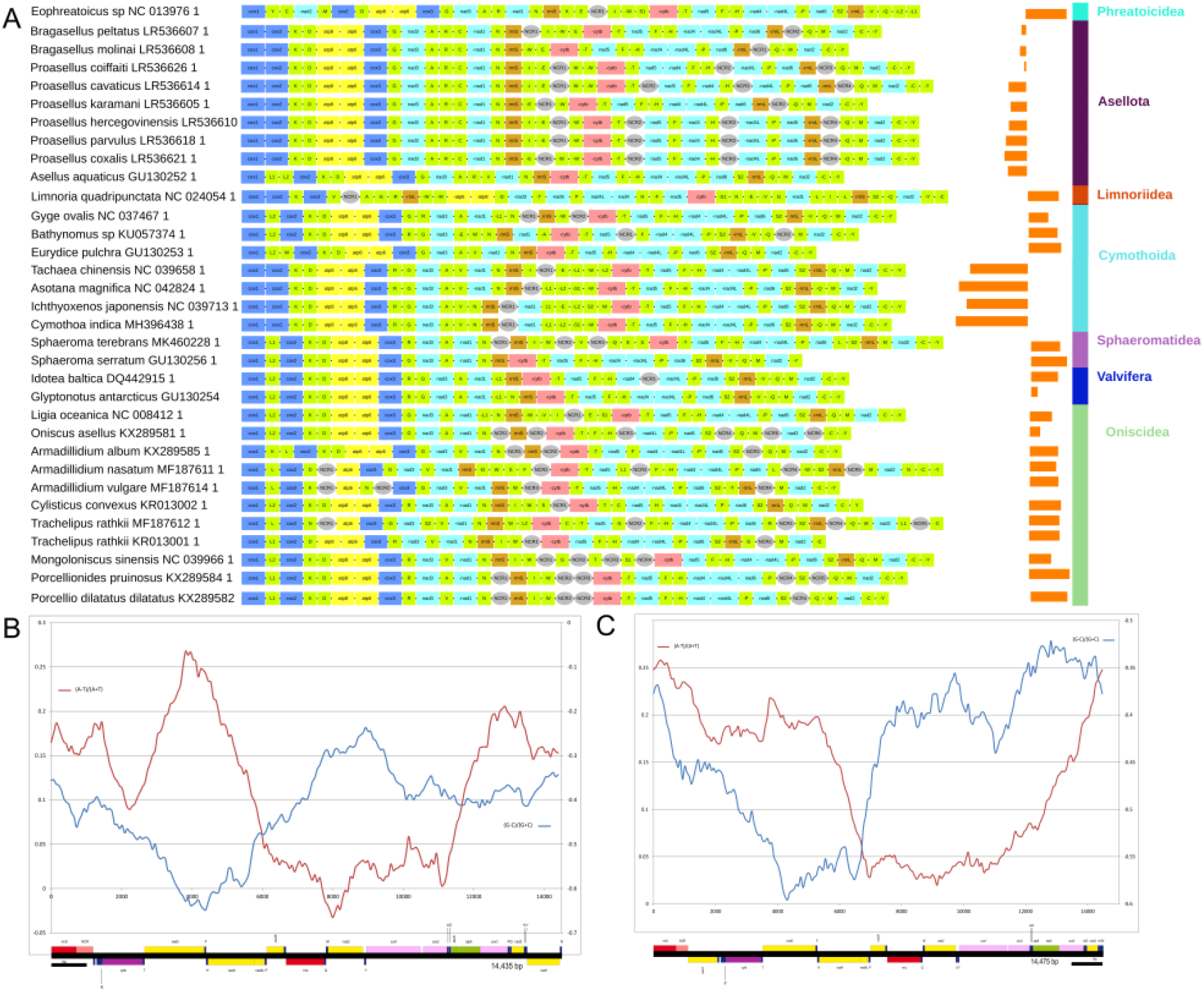
Architecture and skews in the Isopoda. **(A)** Gene orders and skews. (**B)**. Skew plot for *Asotana magnifica*. Global GC Skew = -0.42282, Global AT Skew = 0.11145, WS = 2213, SS = 72. (**C)**. Skew plot for *Cymothoa indica*. Global GC Skew = -0.44197, Global AT Skew = 0. 12874, WS = 3168, SS = 72. See Figure 1 for other details.

**Figure 7.**
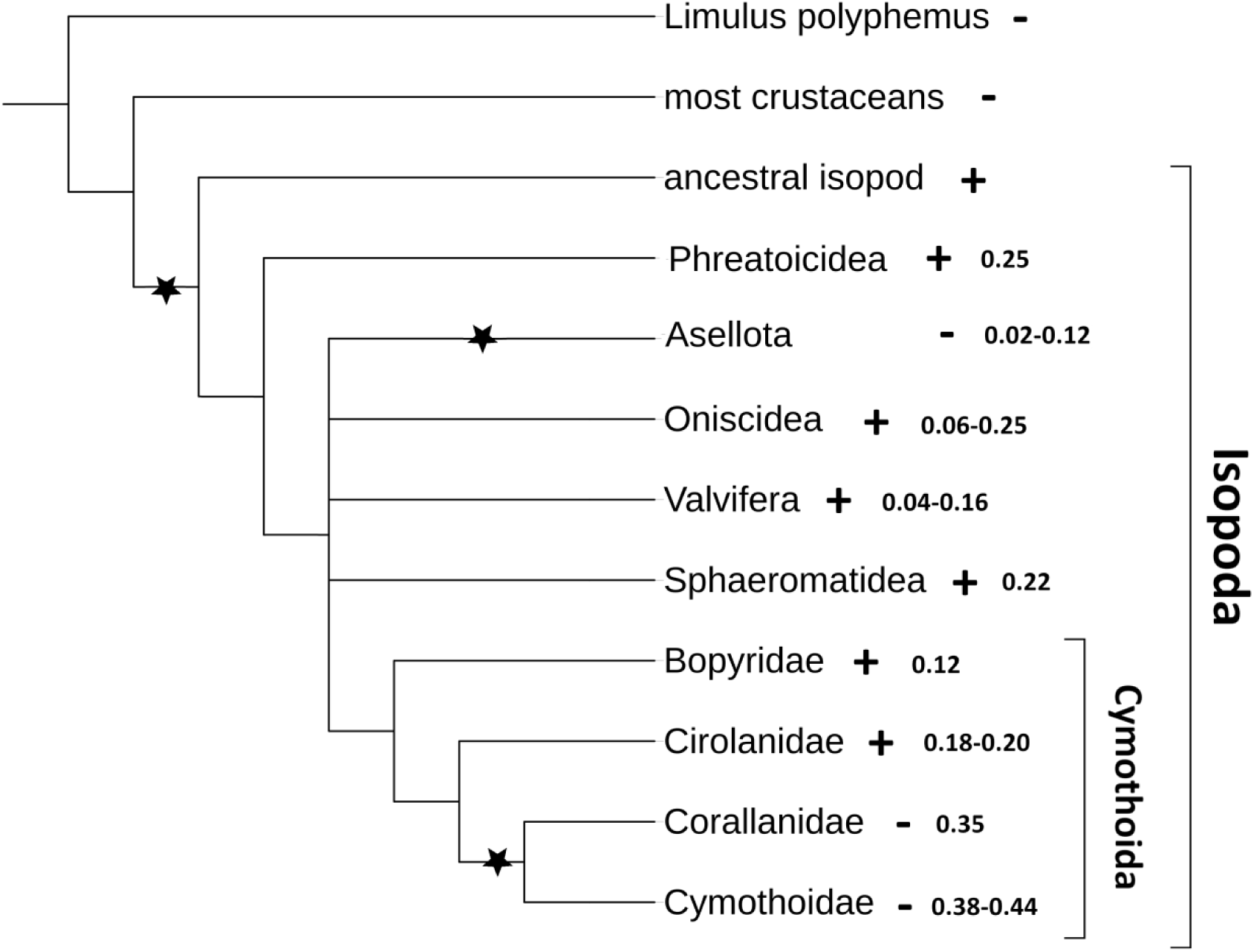
Skews and proposed evolutionary history of the replication origin inversions in the mitogenomes of Isopoda. Isopod suborders are shown at the tips, except for the suborder Cymothoida, where families are shown. +/-indicates whether the GC skew (or direction of the mutational pressure) on the entire majority strand is positive or negative, and the number next to it represents the observed GC skew range in mitogenomes available for the taxon. Star sign indicates a proposed replication origin inversion (ROI) event.

GC skew plot of a basal isopod with a positive overall skew, *Eophreatoicus* sp., exhibits a maximum that fairly well corresponds to the ancestral CR (adjacent to rrnS; Supplementary File S1: Figures S19), which indirectly supports the hypothesis that ancestral isopod ORI can be attributed to a strand switch of the rrnS-CR segment. Two D-I Cymothoidae species, *Asotana magnifica* and *Cymothoa indica* and (GC skews -0.44 and -0.42 respectively), both exhibited GC skew maxima that roughly corresponded to the nad5-F(+)-H-nad4(−) genomic segment (Figure 6B and C). As this segment is relatively well conserved among the isopods (apart from *Limnoria quadripunctata*), it does not explain the ORI. Intriguingly, the D-I species in Cymothoidae and Corallanidae mostly exhibit a highly conserved ancestral arrangement of CR adjacent to the rrnS, and their AT plots do not correspond to the strand distribution of genes, as opposed to most other crustaceans and *Eophreatoicus* sp. In *A. magnifica*, AT and GC skew maxima mirrored each other, but in *C. indica* AT maximum corresponded to the putative CR location (rrnS-NCR). This is indicative of exceptionally high mutational pressures or reduced purifying selection pressures in these two species. Indeed, both overall AT and GC skew values of these two species were among the highest in the entire crustacean dataset (Supplementary File S2). Therefore, although these two species are closely related and exhibit similar overall skews and gene orders, their skew plots are only partially congruent. We suspect that this is more likely to be a consequence of evolutionary recent translocation of nad1 (Fig 6A) than different replication mechanisms, but we cannot reject the latter hypothesis with confidence. In the isopod *Ligia oceanica*, (Kilpert and Podsiadlowski 2006) relied on hairpin structures and flanking TATA and GACT motifs to putatively identify origins of replication. We found that the 476 bp-long major NCR in the mitogenome of *A. magnifica* contains about a dozen regions that may form hairpin structures and several repeated motifs, but this cannot be used to infer the position of the OR as the reverse complement sequence also exhibits similar properties. For example, we identified four repetitions of the TACCCTC motif on the plus strand, which also translates to four repetitions of GAGGGTA on the minus strand, so we cannot say with confidence which strand contains the OR. We also tried aligning the major NCRs of several isopod species, but failed to identify universally conserved segments, thus supporting the observation of (Pie et al. 2008) that CRs in crustaceans tend to be too divergent to be comparable. However, as all Cymothoidae and Corallanidae species sequenced so far exhibit D-I skews (Zou et al. 2020), an ORI event in their common ancestor remains the most parsimonious explanation, so we assume that the rearrangement event behind this ORI comprised only the CR. Sequencing of further mitogenomes from these two and other closely related lineages will be needed to infer the exact phylogenetic timing of this ORI event with confidence.

## Amphipoda (class Malacostraca: subclass Eumalacostraca: superorder Peracarida)

We identified several (putatively) independent ORI events within this order, some of which have been proposed before. Pons et al. hypothesised double-stranded inversion and transposition of a segment comprising *cytb* and *trnS2* to a location adjacent to the putative CR, between *rrnS* and *trnI*, has caused an ORI in the ancestor of the family Metacrangonyctidae (Superfamily Hadzioidea) (Pons et al. 2014). Indeed, species from this family exhibit strongly decreased GC skew magnitude compared to other amphipods (−0.04 vs. -0.27; p=2.86E-10). They have low negative (between -0.05 and -0.01), or even positive (max 0.20 in *Metacrangonyx dhofarensis*), overall skews. However, despite the above rearrangement, the putative CR remained in the ancestral location, adjacent to the *rrnS* (Figure 8). Therefore, given the overwhelming evidence for inverted direction of mutational pressure, we can only hypothesise that the OR underwent strand inversion during the above rearrangement event.

**Figure 8.**
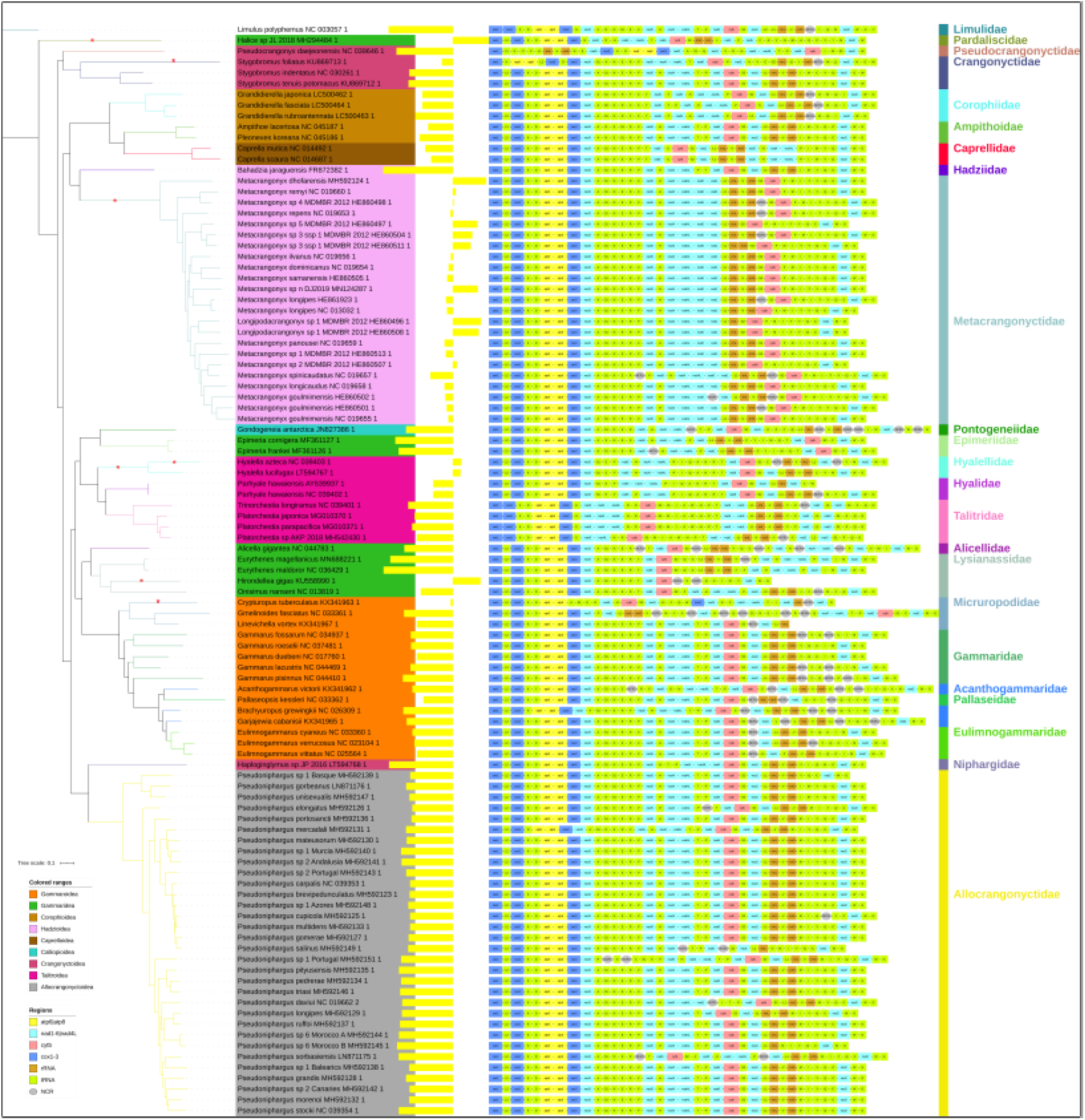
GC skews and gene orders in the Amphipoda.

The overview of skew patterns in all amphipods shows, rather confusingly, similar patterns for most species. For example, comparison of a species with a standard GC skew among the crustaceans, *Bahadzia jaraguensis* (Hadzioidea: Hadziidae) (−0.43), a species from a different superfamily (Gammaroidea: Gammaridae), *Gammarus duebeni* (−0.22), *Metacrangonyx remyi* and *Metacrangonyx ilvanus* (−0.01), reveals that all have very similar skew trends, despite different overall values. All these species exhibit AT skews more or less clearly driven by the strand distribution of genes, and GC skew exhibits a fairly linear decreasing trend (*Metacrangonyx remyi* Figure 9A; Supplementary File S1: Figures S20-26). The difference between the two species with standard skews and the two Metacrangonyctidae species that have GC skews near 0 is that mitogenomes of the latter two species have skews that span a different range of values: from 0.25 to -0.3. In these two species the shift between positive and negative skews corresponds fairly well to the strand distribution of genes. As a similar shift is observable in other mitogenomes, exhibiting only negative GC values, as well, we conclude that the shift is a reflection of mutational constraints imposed by the purifying selection. The best explanation that we can produce for these phenomena is that translocation and strand switch of cytb to a location adjacent to the CR in the common ancestor of Metacrangonyctidae was accompanied by a strand switch and direction inversion of the OR. In this way, the direction of the mutational pressure would change without changing the apparent trend of the skew plot: what used to be the skew minimum (lowest negative value) now became skew maximum. Due to a combination of relatively short evolutionary time (common ancestor of the whole family), low mutational pressure and/or high purifying selection pressure (reflected in the impact that strand distribution has on the skew), these mitogenomes are still in the process of evolving towards positive skews. All this remains hypothetical, and we cannot discount other explanations.

**Figure 9.**
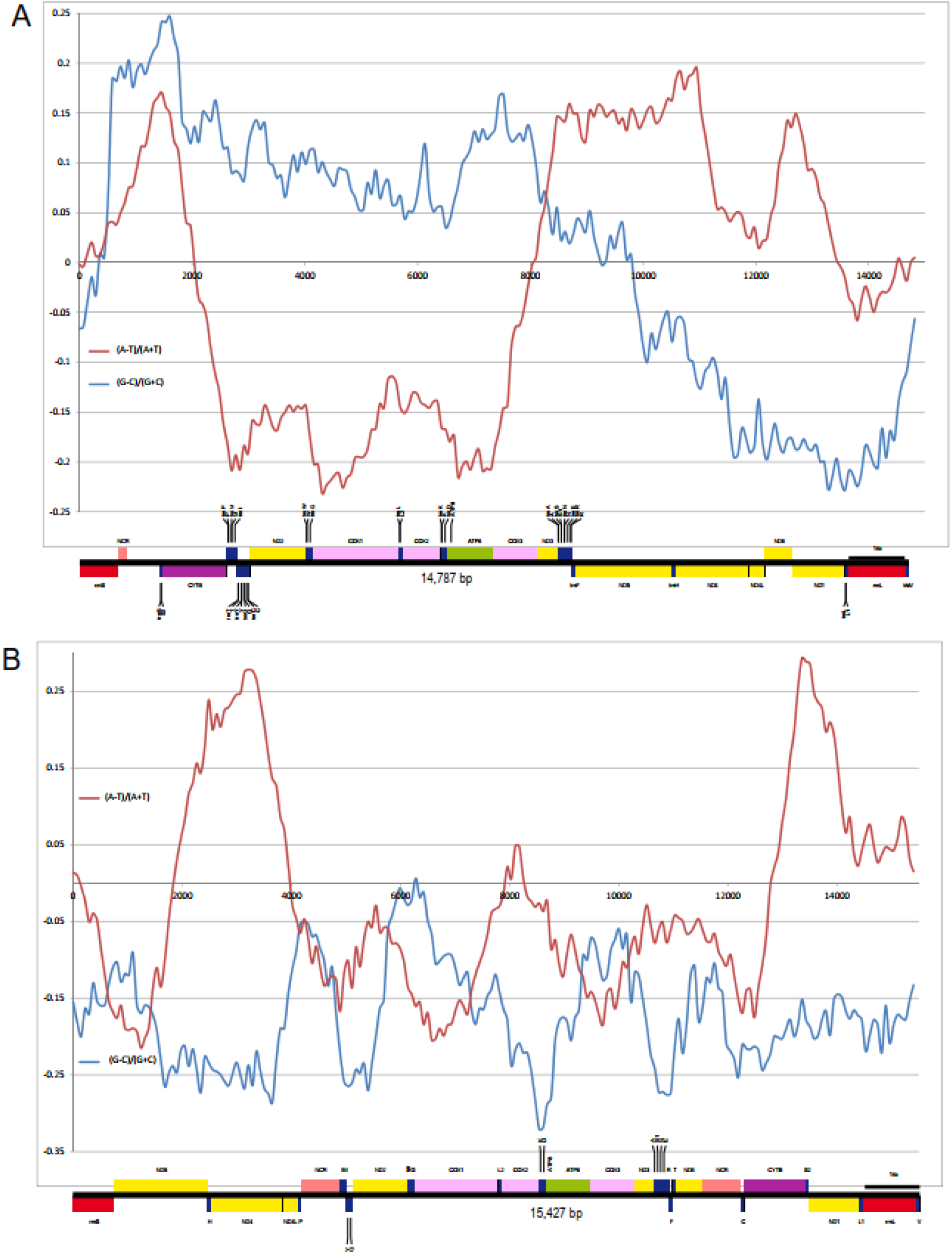
Mitogenomic architecture and cumulative skew plots for selected Amphipoda species. **(A)** *Metacrangonyx remyi*, Global GC Skew = 0.01667, Global AT Skew = -0.01404, Window size = 1238; stepsize = 73. **(B)** *Caprella mutica*, Global GC Skew = -0.17112, Global AT Skew = -0.02280, Window size = 836; stepsize = 73.

The mitogenome of *Caprella mutica* (Caprelloidea: Caprellidae) exhibits a large number of rearrangements and a duplicated NCRs in noncanonical positions, which is also reflected in a noisy GC skew pattern, practically without any trend (Figure 9B). Two species in the superfamily Gammaridea also exhibited positive skews: *Hirondellea gigas* (Lysianassidae) and *Halice* sp. (Pardaliscidae). The skew inversion in *Halice* sp. (0.224) has been observed before (Li et al. 2019), and this ORI can be clearly associated with the translocation, inversion and strand switch rrnL-rrnS (Figure 8). This is the only representative of this family currently available, so we cannot assess the phylogenetic timing of this event. Whereas other Lysianassidae species exhibit standard skews and architecture, *Hirondellea gigas*, which inhabits the deepest point in the ocean (Mariana trench), appears to possess a mitogenome fragmented into two chromosomes (Lan et al. 2016). Both chromosomes have high GC skews (0.17 and 0.54), but closer inspection shows that genes on the second chromosome are all encoded on minus strand in related species. Therefore, this is an annotation artefact, and skew is inverted only on the first chromosome. Although fragmented mitogenomes are common in some metazoan lineages (Nie et al. 2016), including some other arthropods (Rand 2009), this architecture appears to be unique among the crustaceans and deserves a more detailed study.

Both available Hyalellidae (Talitroidea) species also exhibited positive GC skews (≈0.05). *Hyalella azteca* underwent a double-stranded inversion of a large segment comprising *nad1-L1-rrnL-V-rrnS*-CR (−to +), so this appears to be the evidence for the ORI (Figure 8). However, *Hyalella lucifugax* does not exhibit this rearrangement. We also checked the rRNA sequences to corroborate that these are not annotation artefacts. The comparison of GOs of these two species and other related amphipods suggests a rather intriguing, but hypothetical, evolutionary history: an ORI in the common ancestor of all *Hyalella* (or Hyalellidae) species generated by a rearrangement of trnC into the ancestral CR, causing its duplication. This can be observed in the mitogenome of *H. lucifugax* (one of its two CRs is incomplete), but the skew inversion was not observed in the original study (Juan et al. 2016). Subsequently, and probably recently, *H. azteca* (and possibly other species in that lineage) underwent the rearrangement described above (*L1-rrnL-V-rrnS*-CR, - to +). Due to the recency of this event, it is difficult to assess whether this caused an additional ORI from skew patterns (Supplementary File S1: Figures S29-30). It is necessary to sequence more species in this lineage to infer the evolutionary history of GO rearrangements and skew inversions with confidence.

*Stygobromus foliatus* (Crangonyctidae) exhibits a strongly reduced skew (−0.07) compared to the two available congeneric species (≈-0.27). This species exhibits several rearrangements in the order of tRNA genes, including a translocation and strand switch of trnI, and it contains an inverted repeat of the control region (Aunins et al. 2016). On the basis of this evidence, we propose an ORI in this species. *Crypturopus tuberculatus* (Micruropodidae) also exhibits a strongly reduced skew (−0.01) and rearranged architecture compared to the other member of this family, *Gmelinoides fasciatus* (−0.3). Its unique skew pattern was observed before, and the authors proposed an evolutionary recent ORI (Romanova et al. 2016). As the mitogenome is incomplete, we can only tentatively agree with their hypothesis.

## Mysida (class Malacostraca: subclass Eumalacostraca: superorder Peracarida)

Only one species belonging to this order is currently available, *Neomysis japonica*. It exhibits a highly rearranged GO, with most genes on the plus strand, and a low negative overall GC skew (−0.085). trnS-CR segment has been translocated, but trnS remained on the ancestral (minus) strand, and CR increased in size (Figure 10). This large NCR evolves under unique pressures, as its GC skew is positive, whereas the remainder of the genome mostly exhibits negative GC values. This could be interpreted as a sign of a very recent ORI, where only noncoding sequence had time to reach inverted skews due to lower purifying selection pressures, but this is hypothetical. More data are needed for this order.

**Figure 10.**
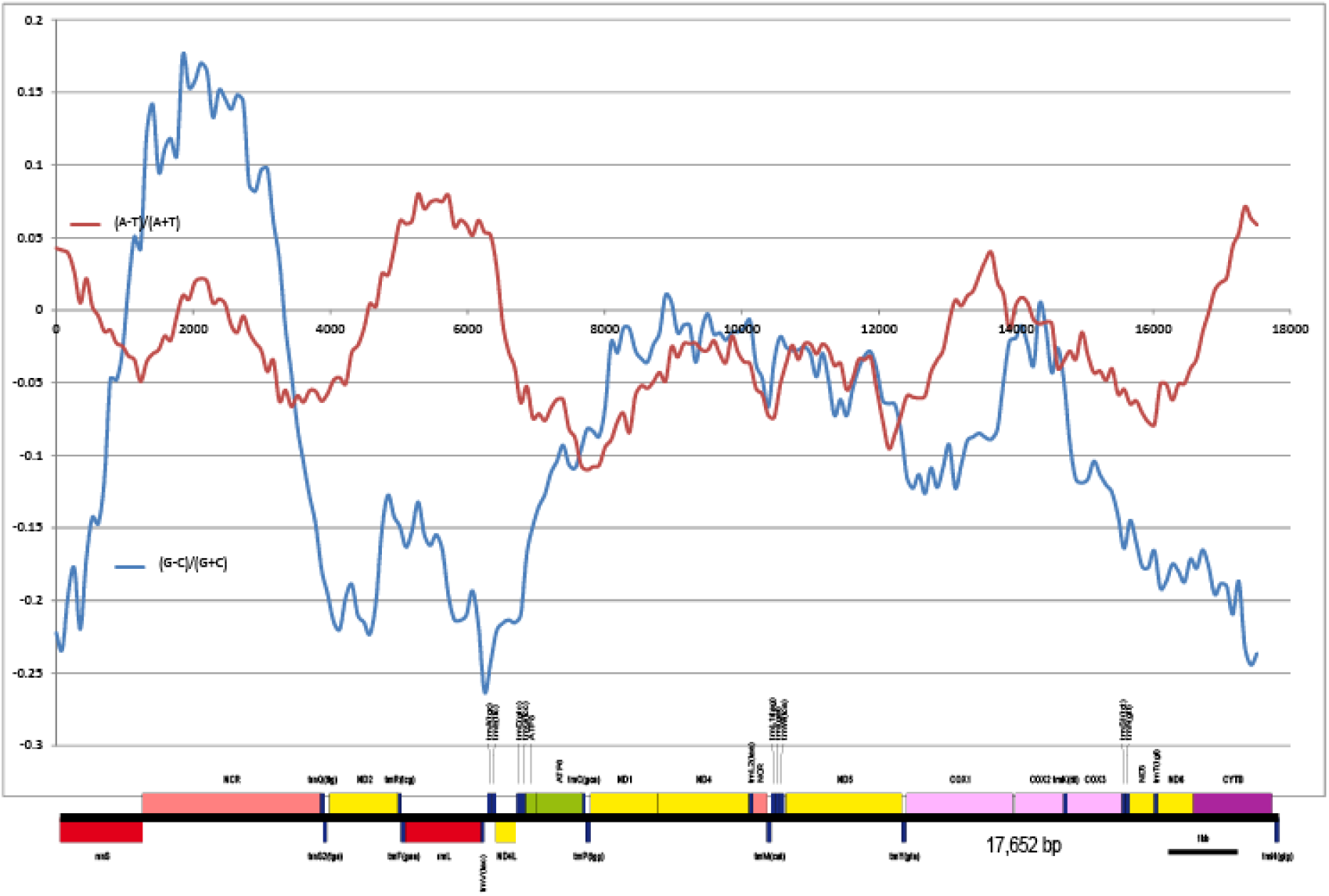
Mitogenomic architecture and cumulative skew plots for *Neomysis japonica*. Global GC Skew = -0.08505, Global AT Skew = -0.02091, Window size = 1500, step size = 88.

## Decapoda (class Malacostraca: subclass Eumalacostraca: superorder Eucarida)

Among the Decapoda, many taxa either possess a highly conserved ancestral crustacean architecture, or rearranged GOs but with conserved strand distribution of the genes associated with the CR in the ancestral GO. In agreement with this conserved GO, they also mostly possess relatively high negative GC skews. A minor outlier was *Belzebub intermedius* (suborder Dendrobranchiata: superfamily Sergestoidea), which exhibited a reduced GC skew magnitude (−0.118) despite possessing a perfectly conserved AAGO. This was the lowest GC skew value among the Decapod species with a conserved AAGO. Also, it did not cluster with the remaining two available species from this family, which also indicates unique evolutionary pressures (or a taxonomic artefact). Among the species with highly conserved AAGO, along with the relatively basal (Shen et al. 2013; Tan et al. 2019) species within the Decapod clade *Penaeus monodon*, discussed previously (Figure 1D), we also plotted skews for a highly derived species *Pachygrapsus marmoratus* (Figure 11). Both GC and AT skew plots exhibited maxima that corresponded to the strand distribution of genes. Primary GC maxima corresponded to nad5-nad4 genes on the minus strand, and secondary roughly corresponded to the nad1-rrnL on the minus strand. As with several other crustacean taxa that exhibited primary GC peak approximately corresponding to these genes, we tentatively assume that this high GC skew is driven by the strand distribution of genes, and not by an OR in this segment of the mitogenome. Below we discuss taxa with inverted skews.

**Figure 11.**
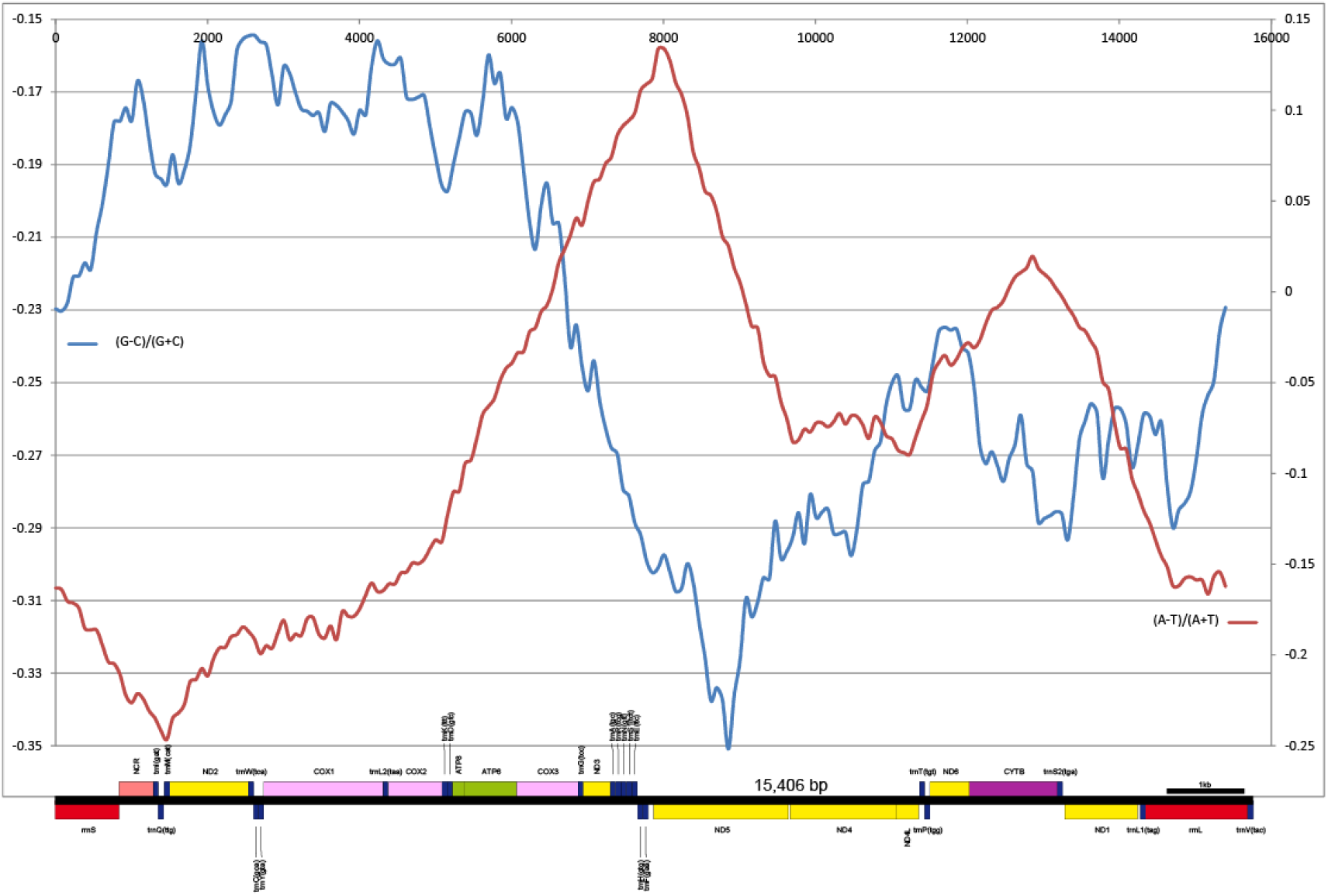
Mitogenomic architecture and cumulative skew plots for *Pachygrapsus marmoratus*. Global GC Skew = -0.23260, Global AT Skew = -0.08173, Window size = 3167, stepsize = 77.

### Astacidea (superfamily)

Inverted GC skews were identified in three out of five families in the superfamily Astacidea: Cambaridae, Astacidae and Nephropidae (Parastacidae and Enoplometopidae had standard skews) (Figure 12). Inverted skews in Cambaridae have been observed before (Kim et al. 2012). All Cambaridae and Astacidae species had positive GC skews, and exhibited a double-stranded inversion of a large fragment comprising 16 genes (approximately *nad5* to *rrnS*). Most species exhibit two large NCRs, one within a large (six genes) tRNA gene cluster upstream from rrnS, and one between the two rRNA genes (rrnS-V-NCR-rrnL) (Figure 12). Within the two families, this architecture is highly conserved, so we can safely infer that after a long period of stasis in mitogenomic architecture evolution (AAGO), their common ancestors underwent a period of highly destabilised, rapidly evolving mitogenomic architecture, and then entered another long period of stasis. This supports the hypothesis of discontinuity in the mitogenomic architecture evolution (Zou et al. 2017). *Cambarus robustus* and *Faxonius rusticus* (Figure 13A and B) exhibited GC skew patterns that support the hypothesis of inverted replication direction, with all values positive and GC maxima corresponding to the ancestral CR.

**Figure 12.**
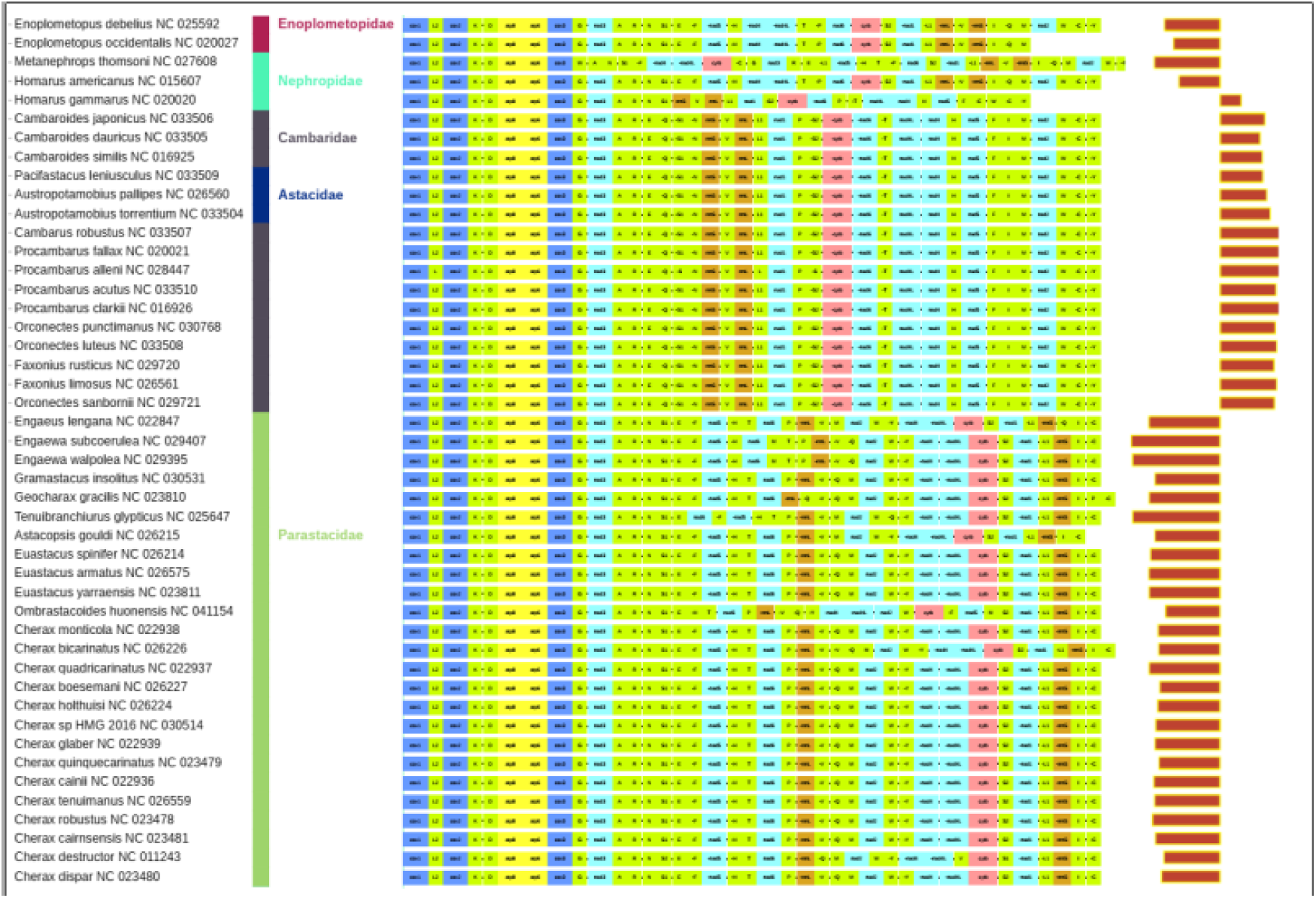
Gene orders and GC skews in the superfamily Astacidea.

**Figure 13.**
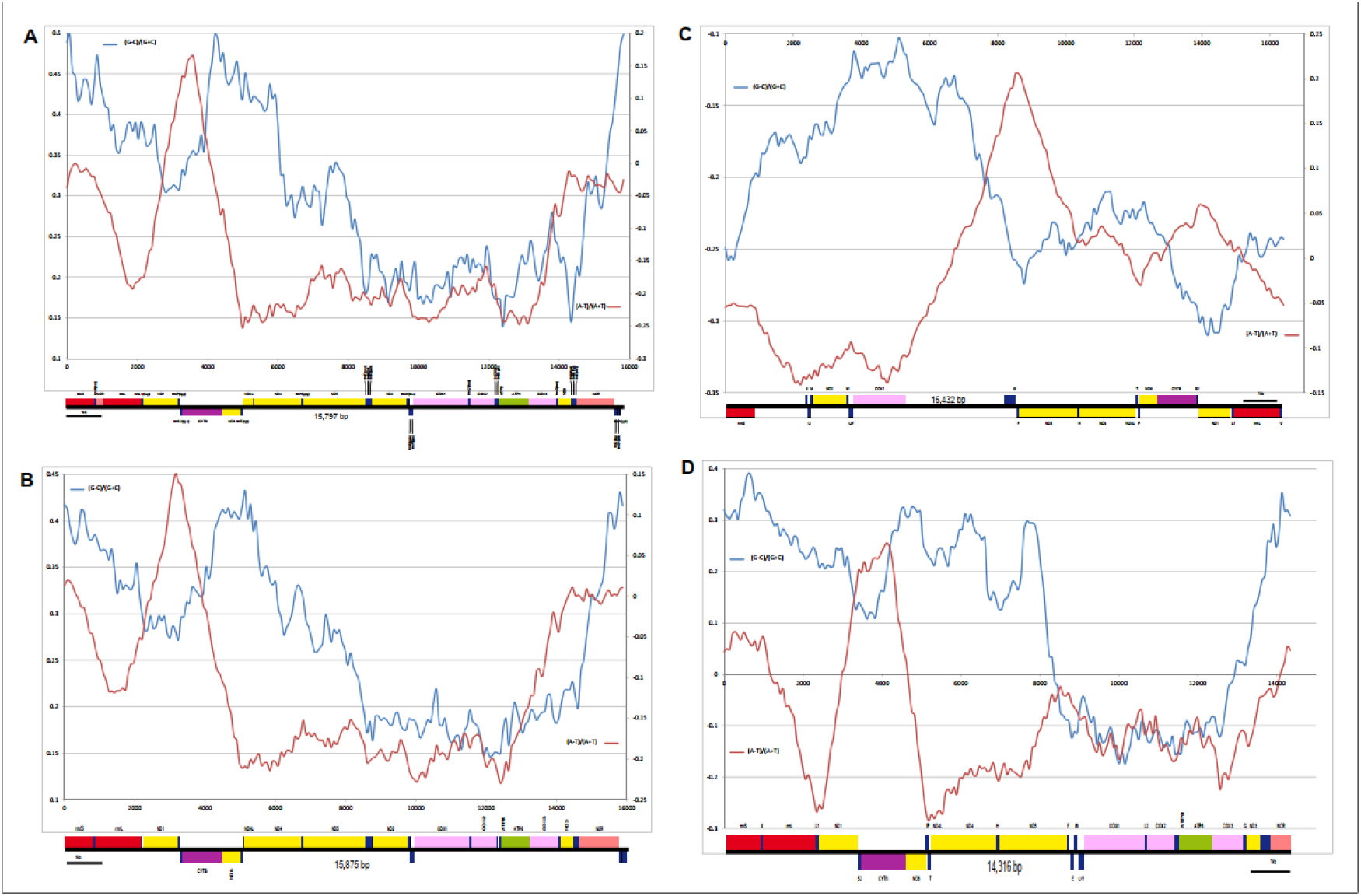
Mitogenomic architecture and cumulative skew plots for selected Astacidea. **(A) *Cambarus robustus*:** Global GC Skew = 0.300, Global AT Skew = -0.134, Window size = 1401; stepsize = 78. **(B) *Faxonius rusticus***: Global GC Skew = 0.270, Global AT Skew = -0.101, Window size = 1851; stepsize = 79. **(C) *Homarus americanus***: Global GC Skew = -0.20431, Global AT Skew = -0.01007, Window size = 3538; stepsize = 82. **(D) *Homarus gammarus***: Global GC Skew = 0.10543, Global AT Skew = -0.08187, Window size = 902; stepsize = 71.

*Homarus gammarus* (Nephropidae) exhibited a unique phenomenon in the entire dataset: two conspecific mitogenomes with very different architecture (Figures 12 and 13). The rearrangements were discussed in detail by the latter of the two studies (Shen et al. 2013; Gan et al. 2019), but neither of the two studies noticed that the first sequenced mitogenome also exhibits a completely inversed (positive) skew. As congeneric *Homarus americanus* exhibits a standard negative skew and architecture relatively similar to the latter sequenced *H. gammarus* mitogenome (negative skew) (Gan et al. 2019), we can safely assume that this is the standard architecture for this species. Intriguingly, this rearrangement between these two mitogenomes is almost identical to the one described for Cambaridae and Astacidae, with the addition of *rrnE. Homarus americanus* (Figure 13C) exhibits a skew pattern comparable that of *P. monodon*, but with inversed primary (nad1) and secondary (nad5) maxima, which suggests the putatively ancestral replication mode, from CR upstream in one segment. GC plot of *H. gammarus* mitogenome with inverted skew exhibits negative values corresponding to the non-inverted part of the genome, and positive values corresponding to the inverted part of the genome (Figure 13D). This indicates that the rearrangement is very recent in evolutionary terms or even that the inversion is an assembly artefact (strand switch), so we shall not treat this is a confirmed ORI. In indirect support of this, (Gan et al. 2019) found that this mitogenome was not fully sequenced. However, as the subsequently sequenced mitogenome (negative skew, MH747083) also exhibits disrupted architecture (duplicated segments) (Gan et al. 2019), we cannot exclude a possibility that this rearrangement is not an artefact, and that it may exist in certain *H. gammarus* lineages. As the first study also failed to identify the *nad2* gene in its mitogenome, this suggests an intriguing possibility that some lineages (or only specimens) may undergo major disruptive mitogenomic events, but continue to be viable for certain periods of time, despite the presumably strongly reduced fitness expected to be produced by the loss of PCGs. We urge sequencing of further mitogenomes from this lineage.

### Anomura (infraorder)

The Anomura exhibit elevated rate of GO rearrangements (Tan et al. 2018), and Coenobitidae exhibited inverted skews (0.14-0.21) (Figure 14). GO rearrangements do not offer a direct explanation for an ORI, and surprisingly the only available species from the sister-family (Schram 2001; Tan et al. 2018) Diogenidae, *Birgus latro*, exhibited an almost identical GO and standard skew (−0.22). The entire clade (both families) exhibits a translocation, inversion and strand switch of *S1-A-nad3-G-L2* (all on minus strand) adjacent to rrnS. We hypothesise that this produced an ORI, and that in *B. latro*, this was followed by a strand switch of OR. Skews were not analysed in the original publication for these mitogenomes (Tan et al. 2018).

**Figure 14.**
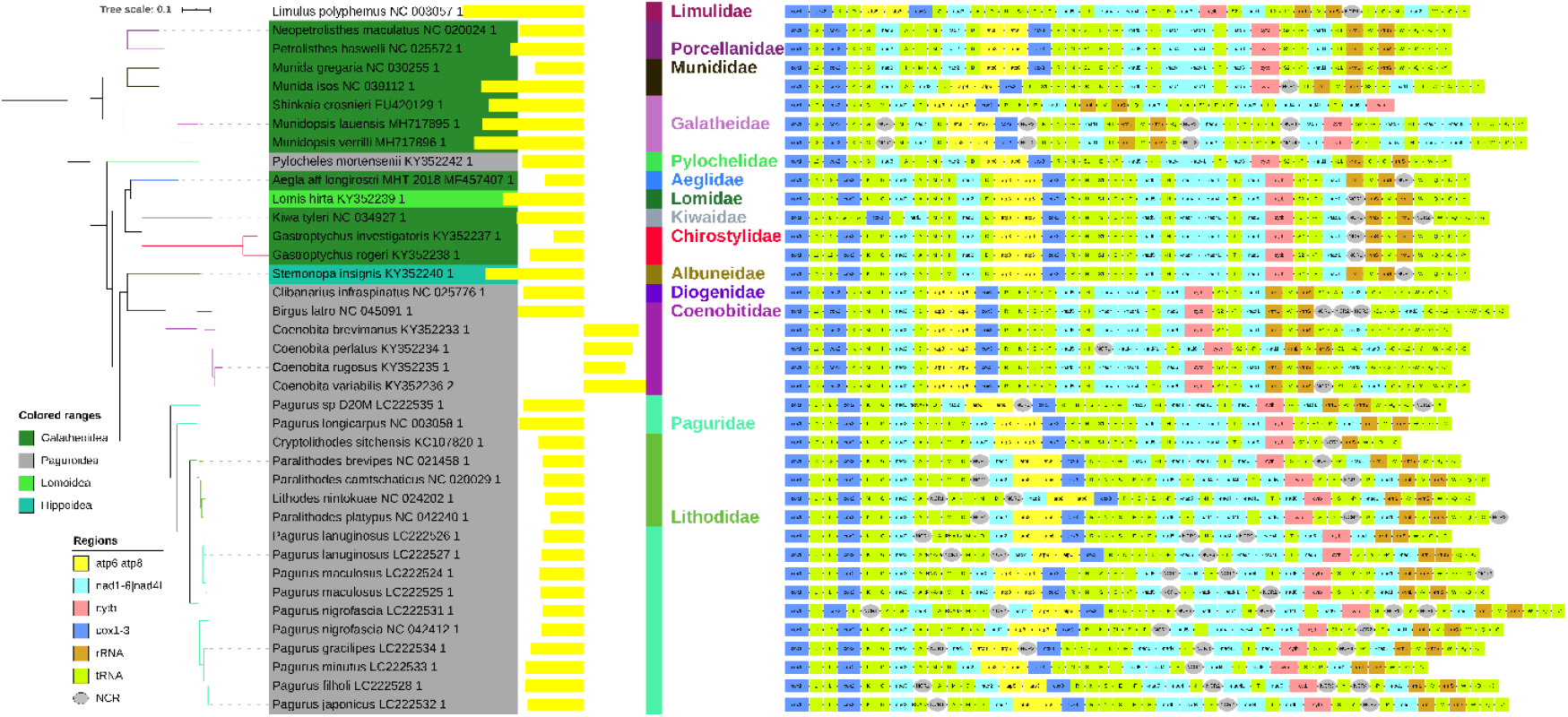
GC skews and gene orders in the infraorder Anomura.

We also identified a rearrangement that suggests an ORI in three families: Chirostylidae, Kiwaidae (superfamily Galatheoidea) and Lomidae (superfamily Lomoidea) (Figure 14). All four available species from these families exhibited almost identical GOs (aside from some tRNA rearrangements in Kiwaidae) and identical double-stranded inversion of the rrnL-V-rrnS segment (Figure 14). However, only *Gastroptychus investigatoris* (Chirostylidae) exhibited a reduced skew magnitude of -0.01, whereas the rest of the species, including the conspecific *G. rogeri* have standard crustacean skews (−0.18 to -0.27). We do not have an explanation why the GC skew magnitude is strongly reduced only in this species.

Another intriguing feature is the superfamily Lomoidea nested within the Galatheoidea. As convergent gene orders are relatively unlikely, the most parsimonious explanation would bean ORI event in the common ancestor of these three families, but the topology inferred here does not support this. This makes us suspect that a mitochondrial introgression event occurred in their evolutionary history. Paguridae and Lithodidae (Anomata) exhibited somewhat reduced skew magnitude, -0.1 and -0.2, but there are no indications of an ORI.

### Axiidea (infraorder)

The Axiidea clade, comprising sister-families (Shen et al. 2013) Strahlaxiidae and Callianassidae, exhibited either strongly reduced negative GC skews (three species) or even positive skews (five species) (Figure 15A). On the basis of the observed GOs and skews, we propose that a most likely scenario includes an ORI in the common ancestor of this entire clade, which resulted in partial or even full skew inversion in most species. The underlying rearrangement can most probably be traced to a double-stranded inversion of the CR-*trnI* (+ to – and inverted direction). The common ancestor of two genera in the Callianassidae family, here represented by *Callianassa ceramica* and *Trypaea australiensis*, appears to have undergone a further autapomorphic ORI (i.e. not shared by other members of the family), caused by a double-stranded inversion of the entire mitogenomic segment *trnD-trnQ-CR-rrnS-rrnL-nad1*. As a result, both species reverted back to negative GC skews. The GC skew plot of *Nihonotrypaea harmandi* (GC=0.08) supports this scenario of inverted direction of replication and mutational pressures, where the upstream segment of the mitogenome exhibits high (positive) skew values, with values slowly decreasing towards zero and finally dipping into the negative zone beginning with cytb (Figure 15B). We speculate that a sharp, narrow trough that corresponds to nad5 is more likely to be associated with stand distribution of genes than with the replication mechanism. From skew profiles, we infer that mitogenomes in this clade are still in the process of skew inversion. This explains why both available *Neaxius spp*. mitogenomes have low negative overall skews (≈-0.06), while exhibiting practically the same GO as *N. harmandi* and other species. Indeed, cumulative skew graphs of these species are almost identical, but shifted ‘lower’ towards the negative values (Figure 15C). This suggests that *Neaxius* mitogenomes are evolving under lesser architecture-driven mutational pressures or under higher purifying selection pressure. From this we infer that the common ancestor of *Callianassa* and *Trypaea* genera probably exhibited a similar GC graph, spanning both positive and negative values. The subsequent ORI once again inverted the replication direction (back to the ancestral upstream direction), so parts of the mitogenome adjacent to the CR are evolving ore rapidly, and thus exhibit negative values, whereas segment distant from the OR is evolving slowly and thus mostly exhibits skews close to zero, including some slightly positive values (Supplementary File S1: Figure S27).

**Figure 15.**
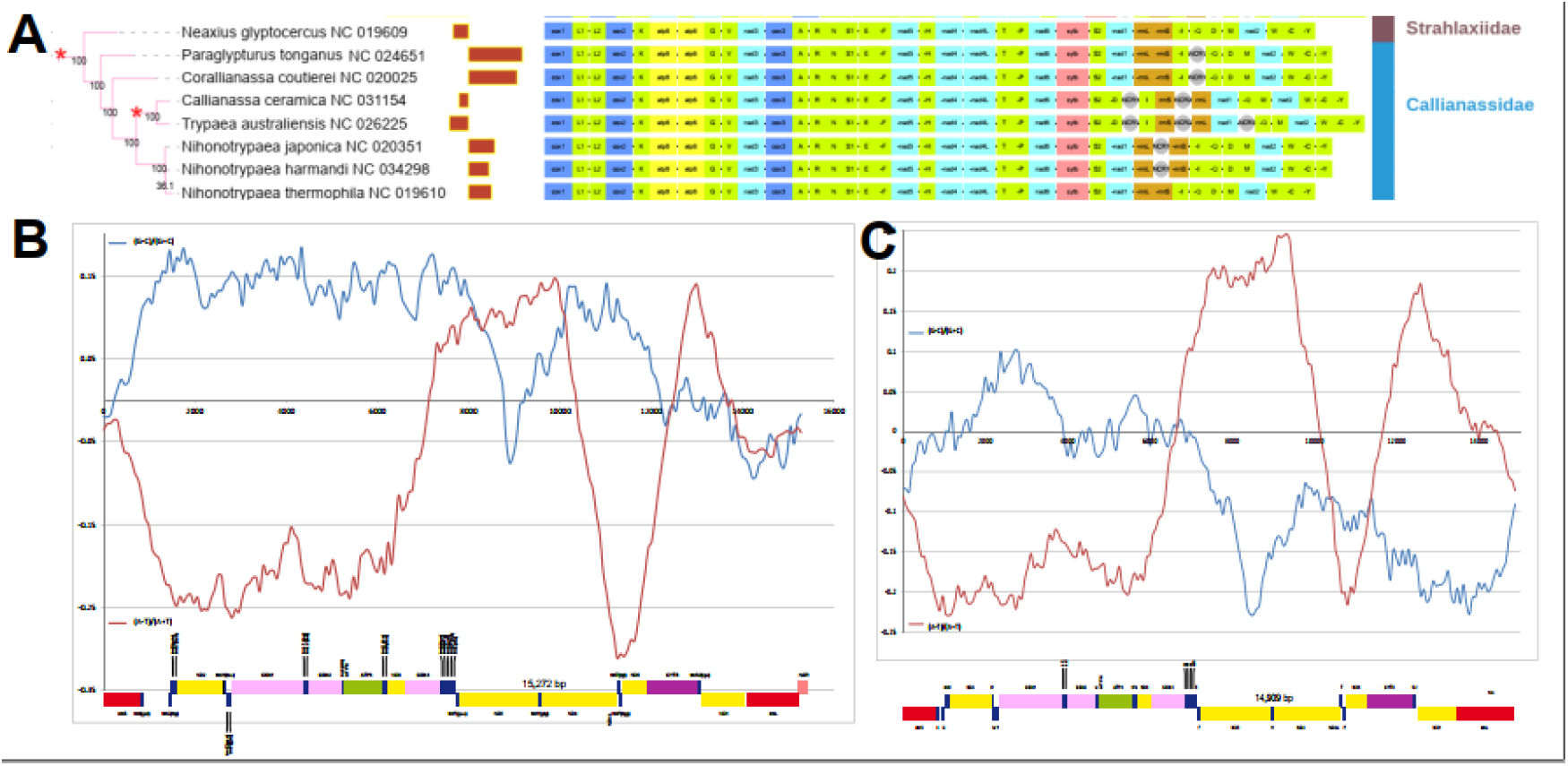
ORI events, gene orders and skew plots in the infraorder Axiidea. **(A)** Putative ORI events are indicated by a star symbol. **(B)** Cumulative skew plot for *Nihonotrypaea harmandi*. Global GC Skew = 0.08431, Global AT Skew = -0.08491, Window size = 1282, stepsize = 76. **(C)** Cumulative skew plot for *Neaxius glyptocercus*. Global GC Skew = -0.06340, Global AT Skew = -0.03313, Window size = 1418, stepsize = 74.

There is an apparent skew inversion in *Austinogebia wuhsienweni* (Upogebiidae), but this is an artefact produced by the authors (unpublished) submitting the minority strand to the GenBank. It may be interesting to observe here that another species in the same family, *Gebiacantha plantae*, exhibits a large number of major rearrangements, comprising the key segment comprising the putative CR. However, the rearrangements did not include strand switches, and skew of this species does not differ from the remaining representatives of this family. This again indirectly supports of the skew inversion model that we relied on in this study.

## Stomatopoda order (class Malacostraca: subclass Hoplocarida)

Species belonging to this order exhibited almost perfectly conserved AAGO and standard skews (Figure 16A), so it was somewhat surprising to observe lesser GC skews (−0.18 on average, p=0.001) and fairly noisy GC skew plots (Figure 16B and C; Supplementary File S1: Figure S28), despite some of the largest optimal window sizes (>3000 bp) inferred herein. All species exhibited approximately mirrored AT and GC maxima, corresponding to a genomic segment (nad5-nad4) encoded on the minus strand. Although the other segment encoded on the minus strand, nad1-rrnL-rrnS, did not produce such strong effects, this suggests that both AT/GC skew patterns are primarily driven by the strand distribution of genes. In conclusion, reduced overall GC skew magnitude and pronounced impacts of strand distribution on GC skew pattern indicate that stomatopod mitogenomes are evolving under weaker architecture-driven mutational pressures and/or stronger purifying selection pressures.

**Figure 16.**
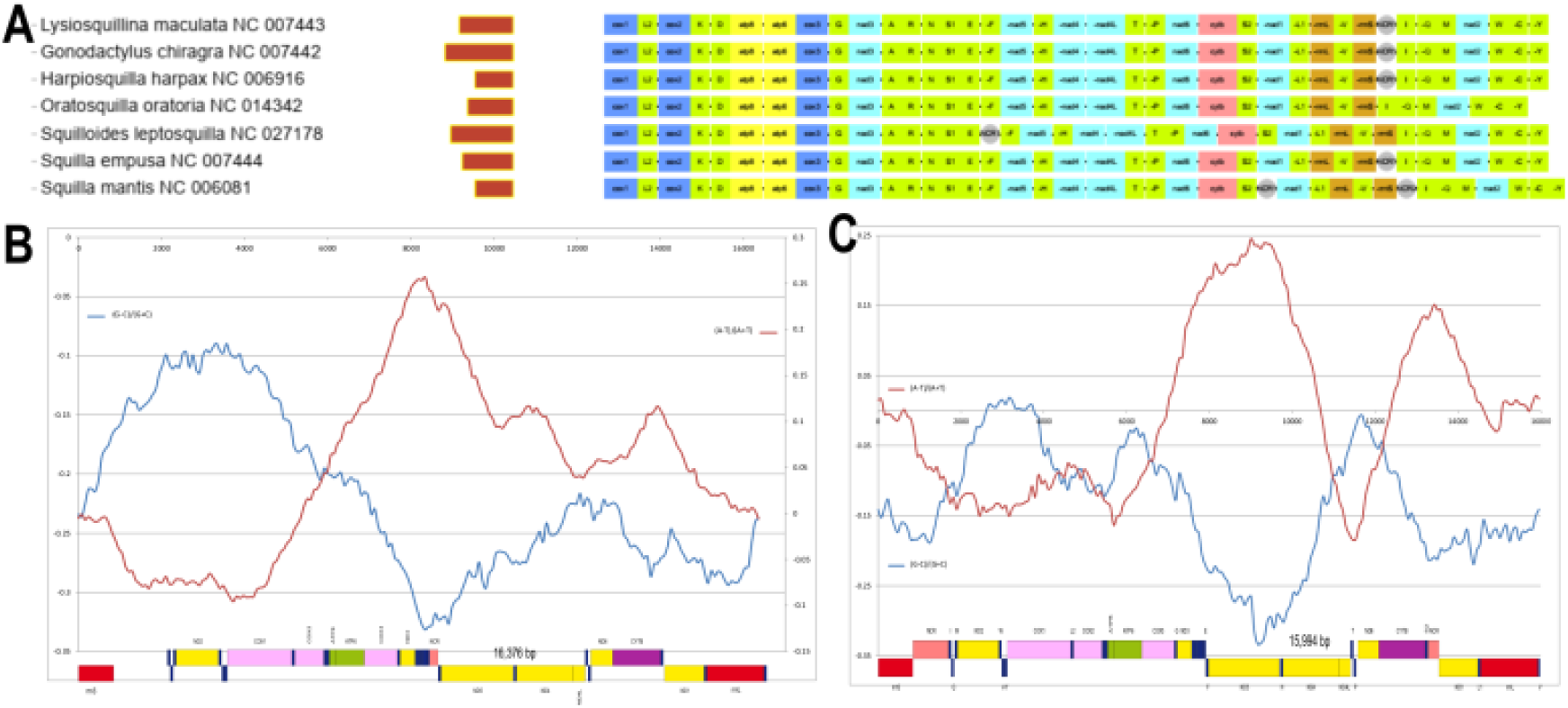
Mitogenomic architecture and skews in the Stomatopoda. **(A)** Gene orders. **(B)** Cumulative skew plot for *Squilloides leptosquilla*. Global GC Skew = -0.21389, Global AT Skew = 0.04891, Window size = 3840, stepsize = 81. **(C)** Cumulative skew plot for *Squilla mantis*. Global GC Skew = -0.13027, Global AT Skew = -0.00062, Window size = 1844, stepsize = 79.

## Discussion

Although some previous studies proposed multiple ORI events in the evolutionary history of crustacean mitogenomes (Hassanin 2006; Pons et al. 2014), none them offered an estimate of the total number. Herein, we inferred 24 putative ORI events (14 of which have not been proposed before): 17 with relative confidence, and 7 speculative (Table 1). Most of these were located at lower taxonomic levels, but there are indications of ORIs that occurred at or above the order-level: Copepoda (Hassanin 2006; Minxiao et al. 2011), Isopoda (Kilpert et al. 2012), and putatively in Branchiopoda (Bellec et al. 2019) and Poecilostomatida+Cyclopoida. Due to a limited number of samples, we cannot discount the possibility that ORIs in Branchiura, Cephalocarida and Rhizocephala may also have occurred at higher taxonomic levels. Our phlyogenetically comprehensive approach allowed us to identify several putative ORI events that did not result in fully inversed skews. Although our dataset is too limited to infer a fully comprehensive evolutionary history of ORI events in the entire crustacean clade, the events not identified by this analysis should be relatively evolutionarily recent (i.e. present only in specific genera and possibly families).

**Table 1.**
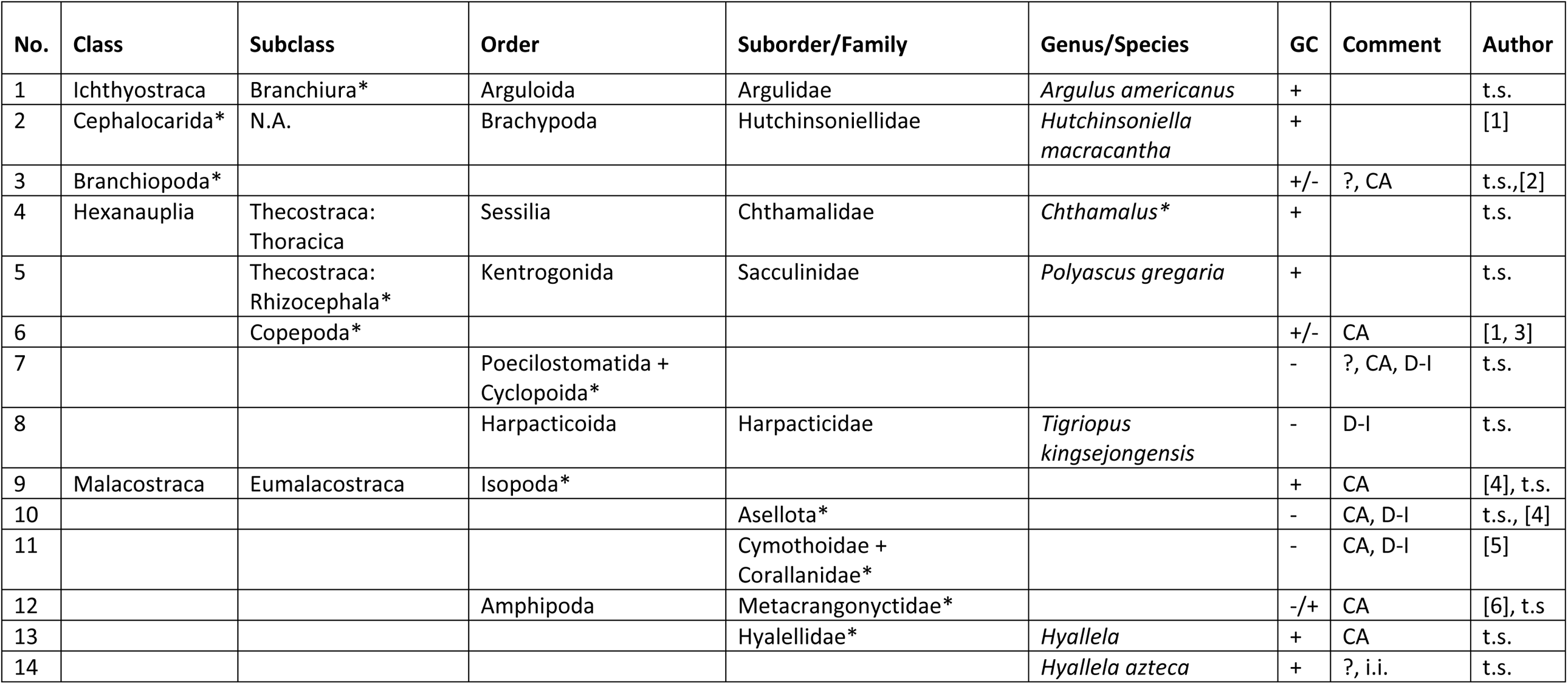
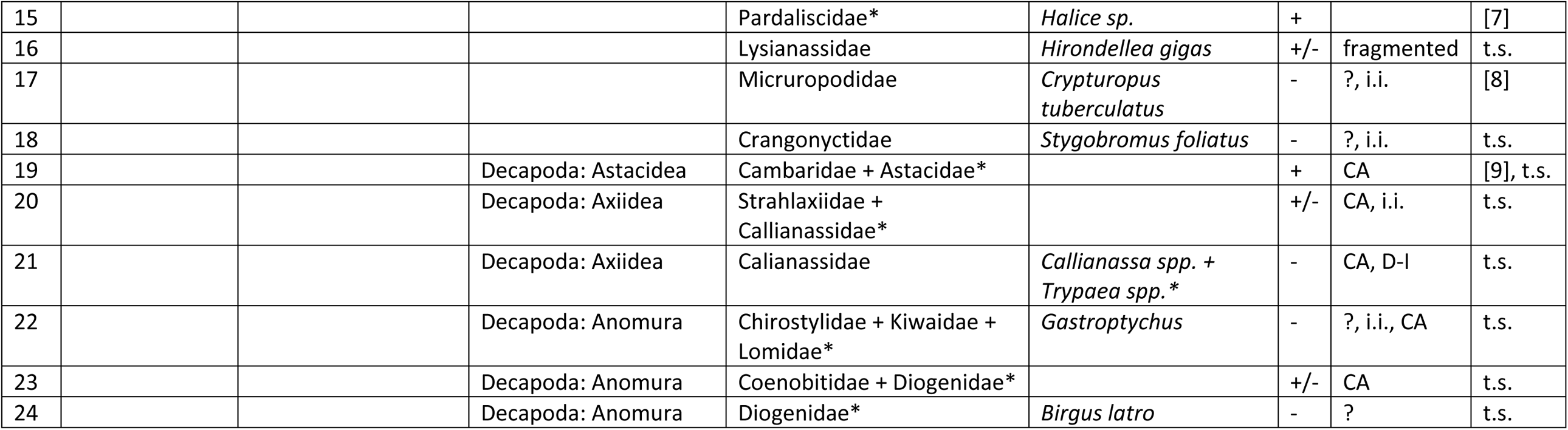
ORI events in crustaceans. * indicates the highest taxonomic level where the ORI could have occurred given the current limitations of our resolution. GC column shows whther the overall skew on the majority strand is positive or ngeative. Comment column: CA = ORI occurred in the common ancestor of the entire taxon; D-I = double-inverted skew (i.e. two ORI events within the same lineage); i.i. = incompletely inverted skew; ? = ORI event that is hypothetical, i.e. not unambiguously supported both by fully inverted skews and mitogenomic rearrangements comprising strand switch of the putative CR. Author column: authors that first observed the inverted skew, where t.s. means ‘this study’; ^#^ The ORI in *Homarus gammarus* (KC107810) is either an assembly artefact or population-specific.

Our analyses also offer an insight into the control of the replication mechanism in crustacean mitogenomes. Despite the discordance between shift points in skew charts and the ancestral location of CR observed in many crustacean lineages, inversions of the mitogenome fragment comprising the ancestral CR (rrnS-CR-trnI) proved to be rather good predictors of skew inversions: most of the observed skew inversions were associated with a strand switch of the genomic segment comprising the ancestral crustacean CR (*rrnS-*CR-*trnI*), and most taxa that underwent a strand inversion of this segment exhibited a change in the skew magnitude compared to the ancestral state. In some lineages, such as isopods, we could not identify rearrangements indicative of ORI, although they clearly exhibited a completely inversed skew. From this we infer that some ORI events can comprise rearrangement events limited to the CR. This was also observed in vertebrates, where some deviations from the default origin of replication were not accompanied by gene rearrangements (Sahyoun et al. 2014). Therefore, regardless of the exact replication mechanism, we can infer with some confidence that there is a close association between the orientation of the ancestral arthropod CR and mutational pressure direction in crustaceans. Our analyses support the previous observation that the evolution of CR is a highly dynamic process in crustaceans, with almost no homology among species at the sequence level (Pie et al. 2008). As a result, in many lineages that exhibited highly rearranged GOs and multiple large NCRs, it was difficult of impossible to infer the location of the CR.

In many lineages, skew plots were not informative for the prediction of replication origin and direction of mutational pressures. This has been reported previously for mitogenomes where the GC skew is not well defined or that underwent recent ORI events, which produce inconsistent, composite signals (Arakawa and Tomita 2007), and in mitogenomes with destabilized mitogenomic architecture and multiple CRs (Sahyoun et al. 2014), as is the case in many crustacean lineages. Intriguingly, in disagreement with a previous report (Xia 2013), we found that shifts in skew plots cannot be used to precisely identify the location of OR even in highly architecturally stable vertebrate (human) mitogenomes. Although skew plots appear to produce ambiguous results with respect to the location of the OR, in some lineages cumulative skew plots produce very clear trends that can be used as an aid to corroborate the ancestral replication mechanism (Lavrov et al. 2000; Saito et al. 2005). Furthermore, they can produce signals meaningful for identification of ORI events that did not result in fully inversed skews or mitogenomes with highly disrupted replication mechanisms.

Intriguingly, although we would expect lineages with conserved ancestral architecture to exhibit similar skew profiles, skew plots of multiple mitogenomes did not behave according to this expectation. This can be explained by multiple factors, but we hypothesise that in most cases it can be attributed to the varying balance of purifying selection vs. mutational pressures among lineages, which can result in varying impacts of strand distribution of genes (discussed in more detail in the next paragraph). The great variability in skew patterns among different crustacean lineages indirectly corroborates the observation that replication mechanism in invertebrates is highly plastic and complex (Lewis et al. 2015; Oliveira et al. 2015). As there is evidence that multiple replication mechanisms may operate simultaneously within some mitochondrial genomes, some of which do not involve a long duration in the single-strand state (Holt et al. 2000; Faith and Pollock 2003; Reyes et al. 2013; Yasukawa and Kang 2018), we cannot exclude a possibility that regulation of replication mechanisms may differ among crustacean lineages with conserved AAGO. Finally, to a lesser degree, skews can be affected by other variables, such as transcription (Touchon et al. 2004; Sahyoun et al. 2014; Fonseca et al. 2014).

We also found that skew plots can be a useful tool to indirectly infer the relative strengths of mutational/purifying pressures in some crustacean lineages: when purifying pressures outweigh mutational, GC skew plots are strongly affected by the strand distribution of genes, and when mutational > purifying, GC skew plots can be even completely (apparently) unaffected by the strand distribution of genes. For AT skews, purifying selection consistently outweighs architecture-driven mutational pressures in crustaceans, so strand distribution of genes fairly well explains the behaviour of the AT skew plot in most crustaceans (and fruit fly), which can be explained by unequal substitution rates between α-bases (A or T) and γ-bases (G and C) (Naylor et al. 1995; Reyes et al. 1998a; Min and Hickey 2007b). However, in some lineages even the GC skew plots corresponded to the strand distribution of genes, which implies that these lineages are evolving either under elevated purifying selection, or under reduced architecture-driven mutational pressures. This observation has very important repercussions for phylogenetic and evolutionary studies, as it implies that not only the relatively rare ORI events, but also gene strand switches, which are very common in some lineages (such as isopods), can produce very strong compositional biases, which in turn may affect phylogenetic and evolutionary analyses.

In more detail, mitogenomes of crustacean taxa with conserved ancestral architecture have been evolving under unidirectional architecture-driven mutational pressures for approximately 400 million years (Lavrov et al. 2000), so they have long reached mutational saturation with respect to synonymous mutations. As a result, their evolution is a product of two asymmetrical evolutionary pressures: architecture-driven mutational pressures and purifying selection. In a stable environment, this is expected to produce very slow evolutionary rates. An ORI event changes the direction of nonadaptive mutational pressures, thus making thousands of sites in a mitochondrial genome available for synonymous mutations, which can produce apparent evolutionary bursts. On a smaller scale, similar bursts occur when a gene that has reached mutational saturation undergoes a strand switch. This was observed previously in bacterial genomes, and the authors proposed that different strand-specific substitution models should be used along different branches in a phylogenetic analysis of species that underwent strand switches of genes (Marín and Xia 2008). The authors proposed that a similar strand-biased model may also be needed in modelling mitochondrial genomic evolution (Marín and Xia 2008). Finally, even a same-strand relocation of a gene may produce notable changes in mutational pressures (for example from a location far from the OR to a location adjacent to the OR). Therefore, most types of mitochondrial architecture rearrangements are expected to produce varying levels of compositional heterogeneity. Similarly, it has been proposed that the key factor underlying the species differences in bacterial genomes is not the functional environment of the proteins but rather the location of the genes encoding those proteins (Min and Hickey 2007a). Importantly, rearrangement events may produce evolutionary bursts even at the protein level, as some amino acids can be relatively easily replaced with amino acids with similar properties (Min and Hickey 2007a; Botero-Castro et al. 2018).

In indirect support of this, our phylogenetic analyses produced highly unorthodox crustacean topologies, with numerous LBA artefacts, regardless of the fact that we used amino acid datasets and an algorithm designed to account for compositional heterogeneity. This supports our previous observation that composition skews in some crustacean lineages are too strong to be resolved using the currently available strategies for compositional heterogeneity (Zhang et al. 2019b; Zou et al. 2020), and indirectly explains a proportion of the contradictory hypotheses put forward for the phylogeny and taxonomy of crustaceans in the past (Timm and Bracken-Grissom 2015; Lozano-Fernandez et al. 2019).

Importantly, apart from producing noise in the phylogenetic analyses, these compositional biases can produce misleading signals in other types of evolutionary analyses as well, such as dN/dS ratios, codon usage bias, base composition, branch length comparison, etc. We suspect that ORI-driven and gene strand switch-driven evolutionary bursts were often mistakenly misinterpreted as evidence for rapid adaptive evolution in a lineage. For example, (Tan et al. 2017) have observed that Axiidea exhibit a different codon usage pattern and faster evolution than closely related taxa, and they speculated that this may be associated with their burrowing lifestyle. Although we do not reject a possibility that their unique life history played a role in their mitochondrial evolution, we suspect that unique codon usage pattern and fast evolution may be largely attributable to the ORI events that we identified in this clade. In another example, a study (Vanschoenwinkel et al. 2012) found that the fastest rates of *cox1* evolution (4.9% mya^−1^) in invertebrates have been reported in barnacles (Wares 2001). This is in clear disagreement with our findings that most of the barnacles evolve at exceptionally low mutational pressures for the crustacean dataset. However, this rate was inferred on a dataset comprised of *Chthamalus* species (Wares 2001), for which we inferred an ORI event in this study. Therefore, we strongly suspect that the elevated evolutionary rate is largely a product of the ORI event, and not reflective of the average evolutionary rate in other barnacles. Finally, the fact that some, but not all, architectural rearrangements will produce mutational bursts may also be a partial explanation for the observation that there is only a weak association between the GO rearrangement and base substitution rates in mitogenomes of many animal lineages (Shao et al. 2003; Hassanin 2006; Marlétaz et al. 2017; Tan et al. 2019). Therefore, in agreement with previous observation that demonstrating the occurrence of adaptive substitutions in mtDNA evolution can be difficult (Lavrov and Pett 2016), all studies aiming to study the evolution of mtDNA should pay close attention to architectural rearrangements and misleading mutational signals that they may produce.

These findings do not imply that mitogenomes should not be used for evolutionary studies in crustaceans. More than 40 decapod families in our dataset exhibited standard crustacean skews, and even a handful of orders, such as Stomatopoda, do not exhibit any ORI events. Due to their comparative advantages over other molecular markers, including unilinear inheritance, lack of recombination, higher resolution than single-gene markers, and rapidly growing availability, mitogenomic data can still be a very valuable tool for inferring lower-level (e.g. superfamily and family) phylogenies in all crustacean lineages that do not exhibit skew inversions. Additionally, this stark contrast between these highly conserved decapod lineages and disrupted architecture of Copepoda, Isopoda and some other lineages supports the hypothesis of high discontinuity in the evolution of mitochondrial architecture between different lineages (Zou et al. 2017). As the evolutionary forces behind this discontinuity remain almost completely unresolved, crustaceans may represent a good model to study this puzzling question.

## Abbreviations

CR: control region,
RO: replication of origin,
ROI: inversion of the replication of origin,
D-I skew: double-inverted skew,
LBA: long-branch attraction

