## Supplementary File S1 for "Evolutionary history of inversions in the direction of architecture-driven mutational pressures in crustacean mitochondrial genomes"

**Supplementary File S1. Cumulative skew plots for selected crustacean mitogenomes and calibration.**

In cumulative skew plot charts, Y-axis shows the skew magnitude and X-axis the mitochondrial sequence (number of bases). Some figures are shown with the corresponding mitochondrial genome map, where OH and OL indicate approximate locations of the origins of replication of two mitochondrial strands (if known). WS = window size, SS = step size. Average values for each window are shown at the start point, unless if indicated otherwise in the figure. GC and AT indicate which of the correspnding two skews is shown: AT skew = (A-T)/(A+T), GC skew = (G-C)/(G+C).

### Calibration

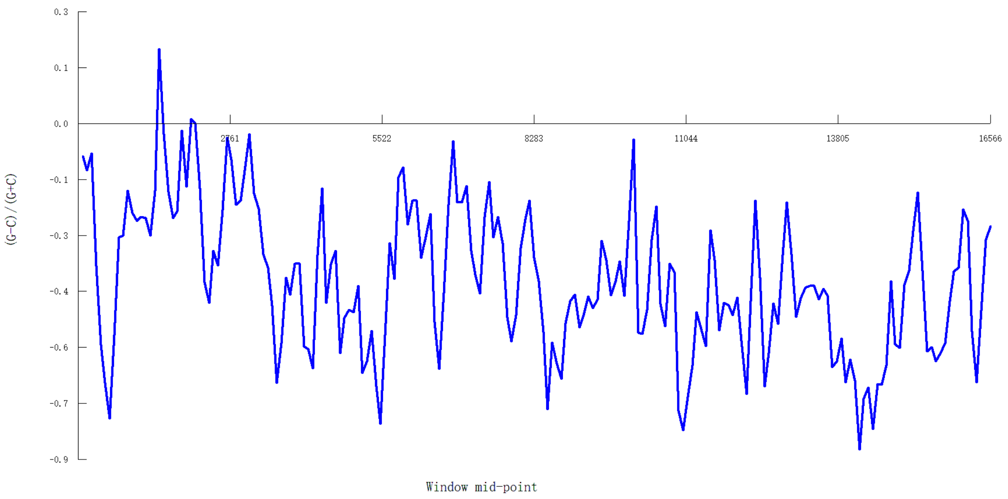


Figure S1. GC plot for *Homo sapiens* (NC_012920) with manually set WS to 165 (SS = 82).

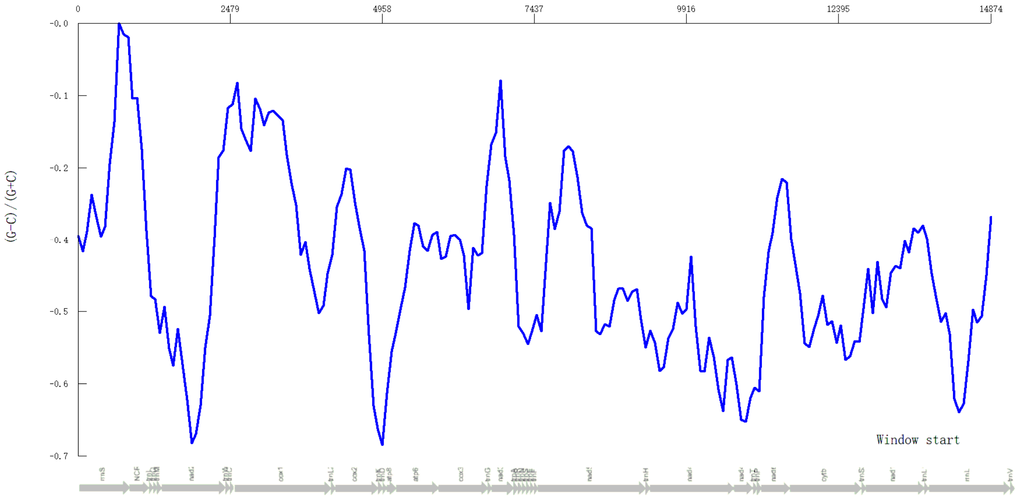

Figure S2. *Limulus polyphemus* GC skew plot with manually set WS to 500 (SS = 74).

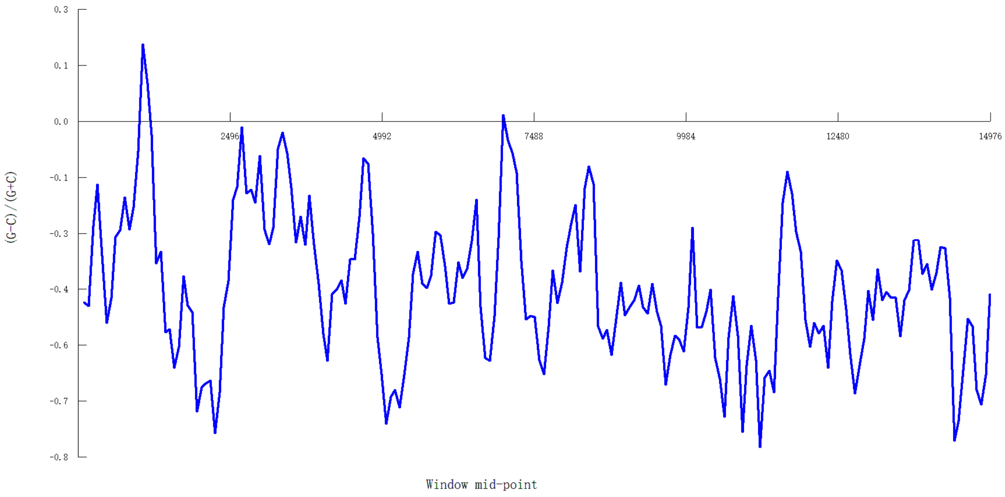


Figure S3. *Limulus polyphemus* GC skew plot with manually set WS to 200 (SS = 74).

### Ichthyostraca

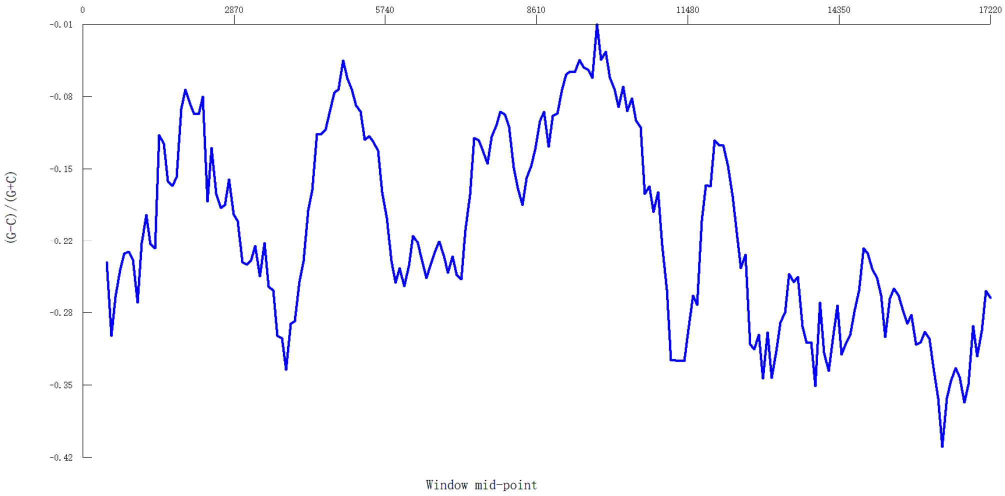


Figure S4. *Cypridopsis vidua* GC (WS = 912, SS = 83).

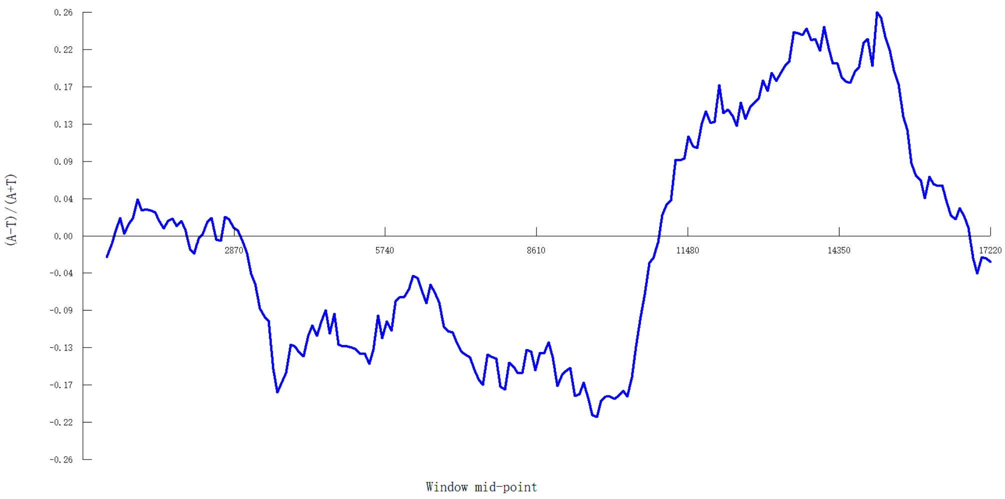


Figure S5. *Cypridopsis vidua* AT (WS = 912, SS = 83).

### Branchiopoda

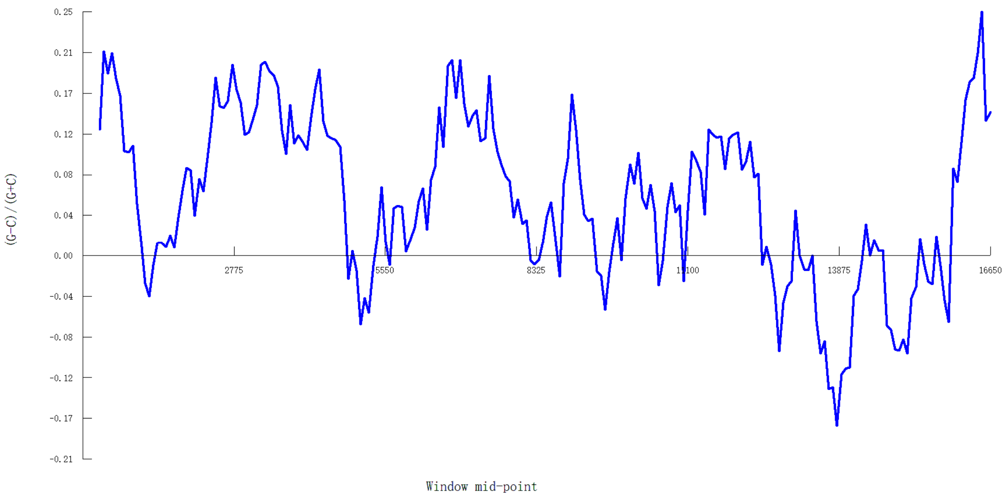


Figure S6. *Diaphanostoma dubium* GC (WS = 620, SS = 76).

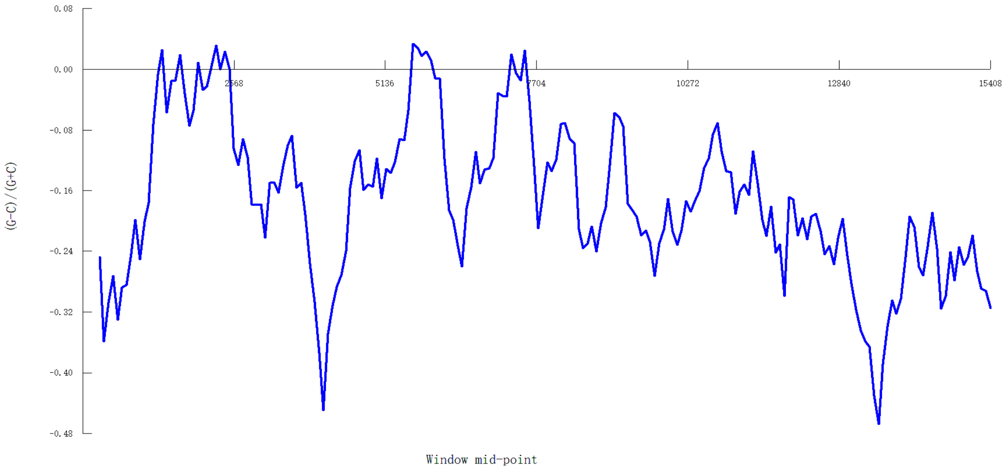


Figure S7. *Triops cancriformis* GC.

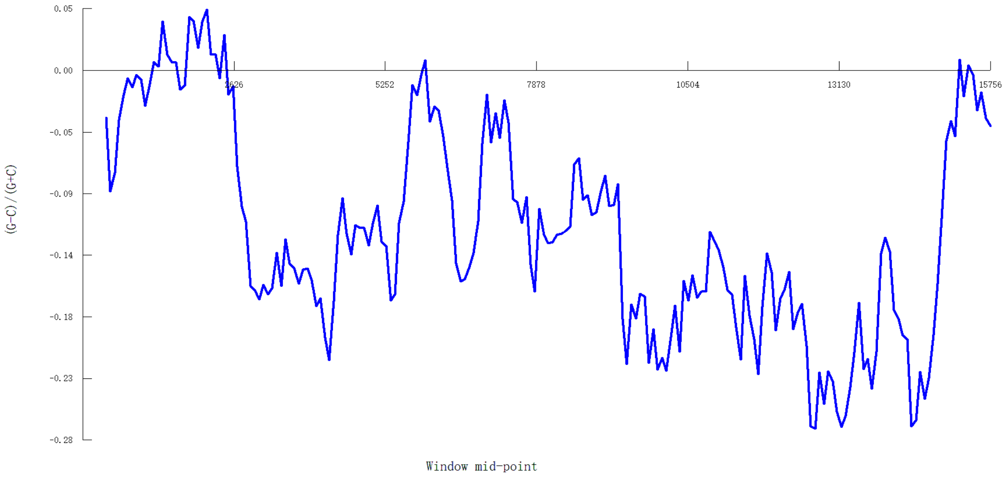


Figure S8. *Daphnia pulex* GC.

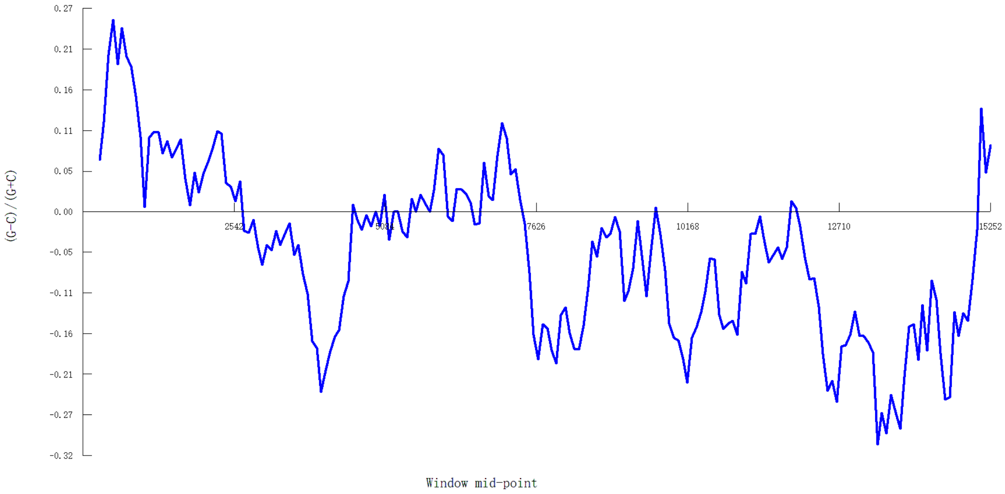


Figure S9. *Daphnia magna* GC.

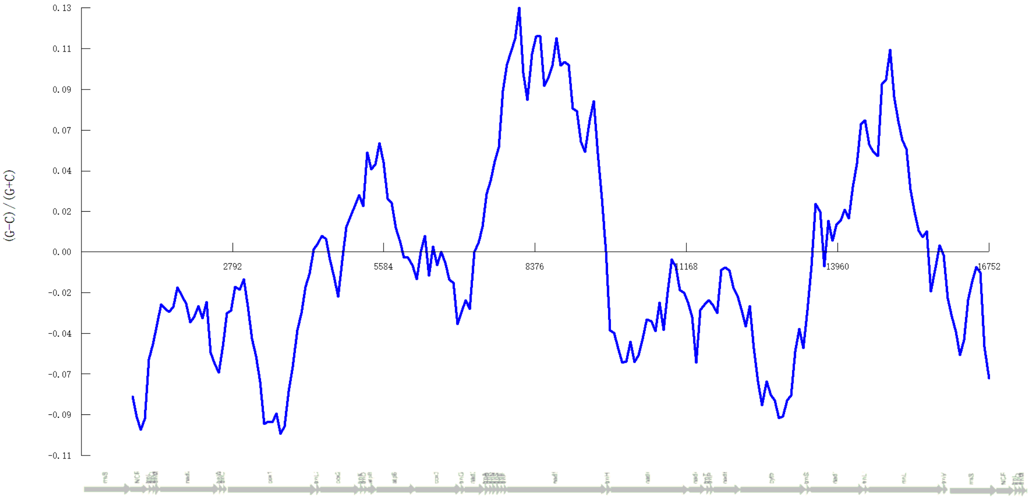


Figure S10. *Artemia franciscana* GC.

### Copepoda

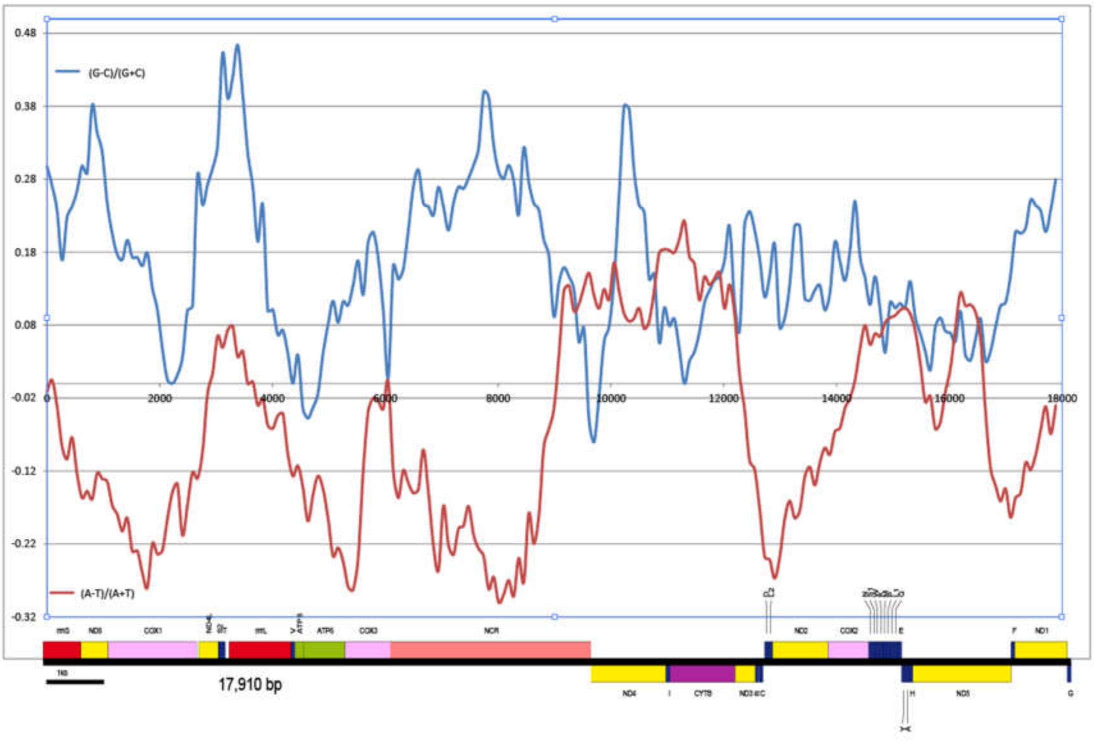


Figure S11. *Calanus hyperboreus.* Automatic WS = 560; SS = 89. Global GC Skew = 0.15871, Global AT Skew = -0.06013.

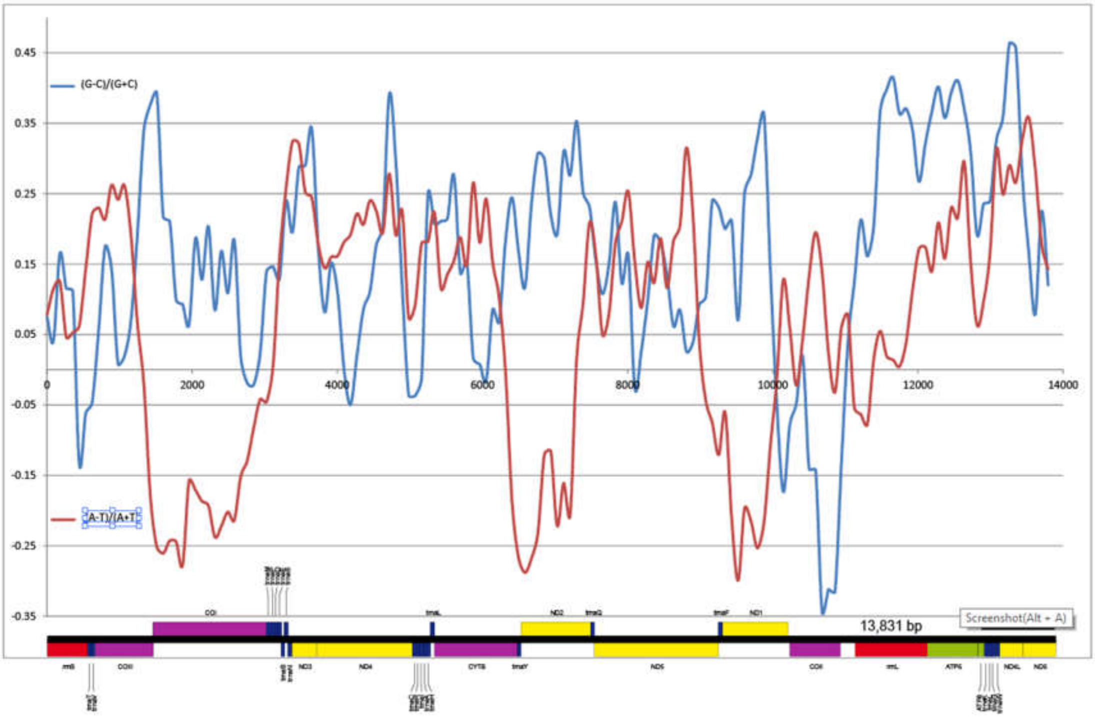


Figure S12. *Amphiascoides atopus*. WS = 299; SS = 89. Global GC Skew = 0.14545, Global AT Skew = 0.06745.

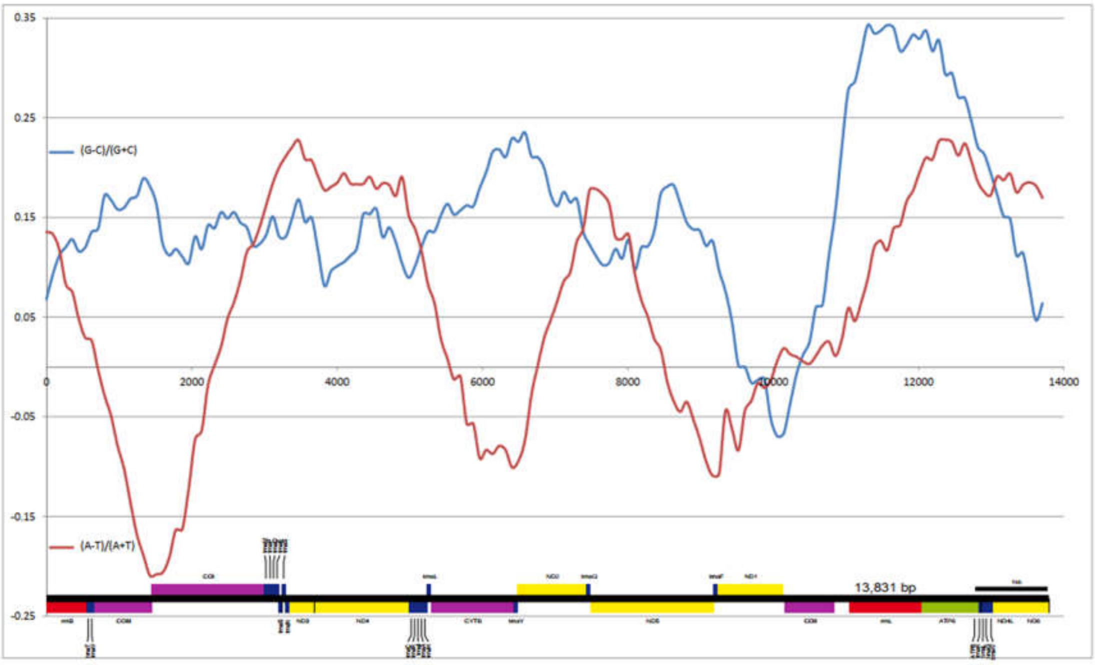


Figure S13. *Amphiascoides atopus.* WS = manually set at 1500; stepsize = 89.Global GC Skew = 0.14545, Global AT Skew = 0.06745,

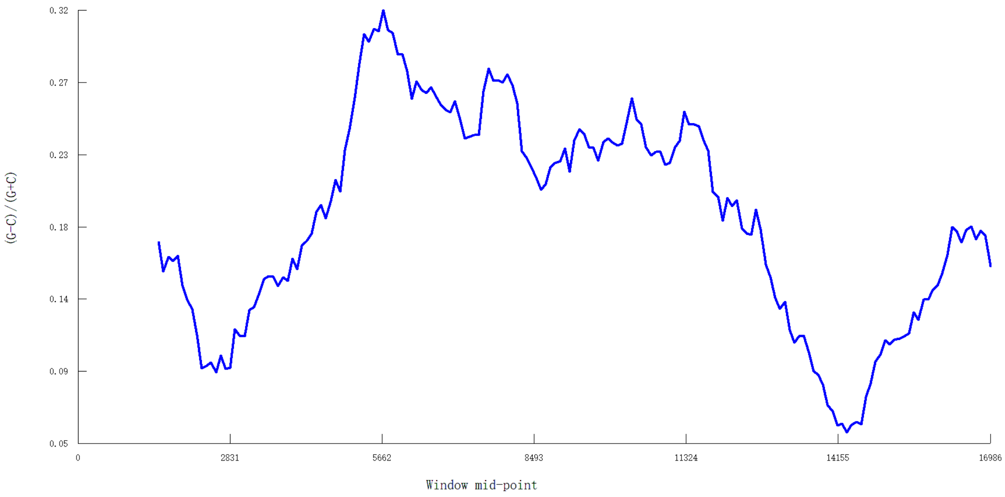


Figure S14. *Lepeophtheirus salmonis* GC.

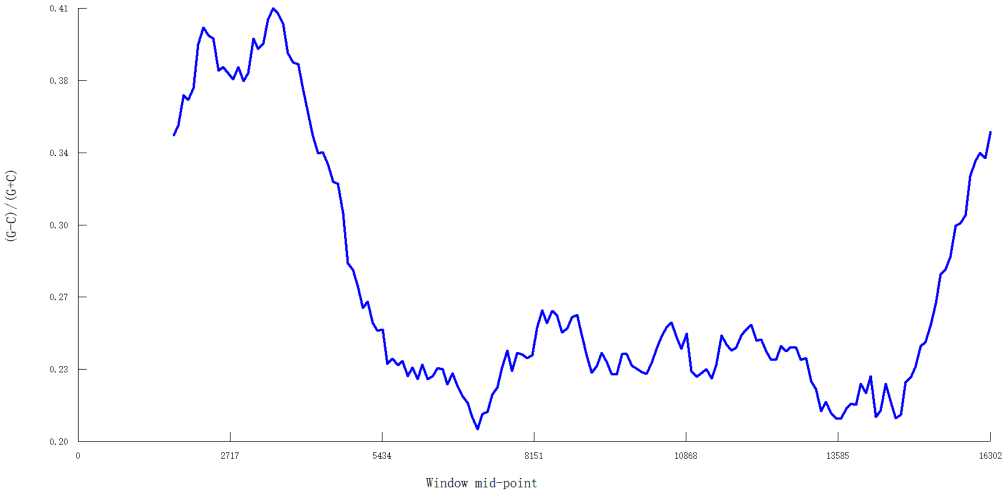


Figure S15. *Tigriopus californicus* GC.

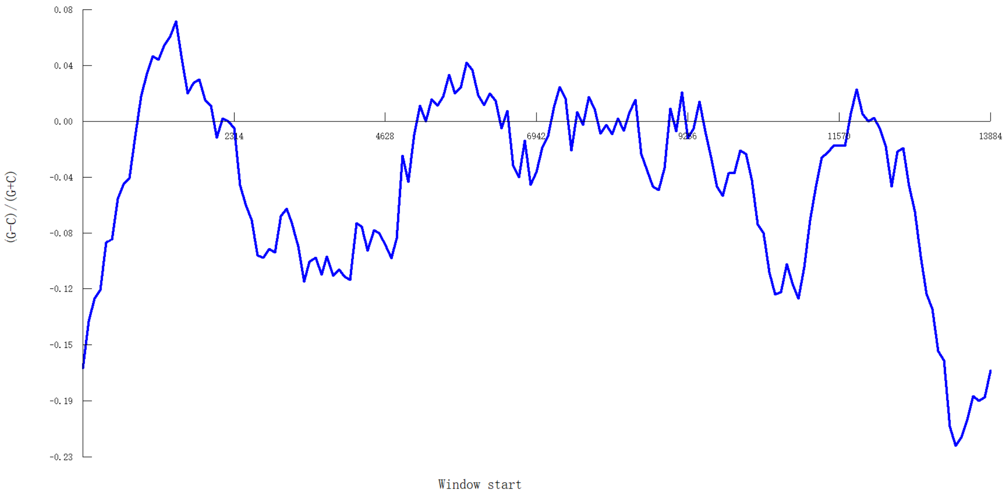

Figure S16. *Sinergasilus polycolpus* GC (manually to 1500 bp)

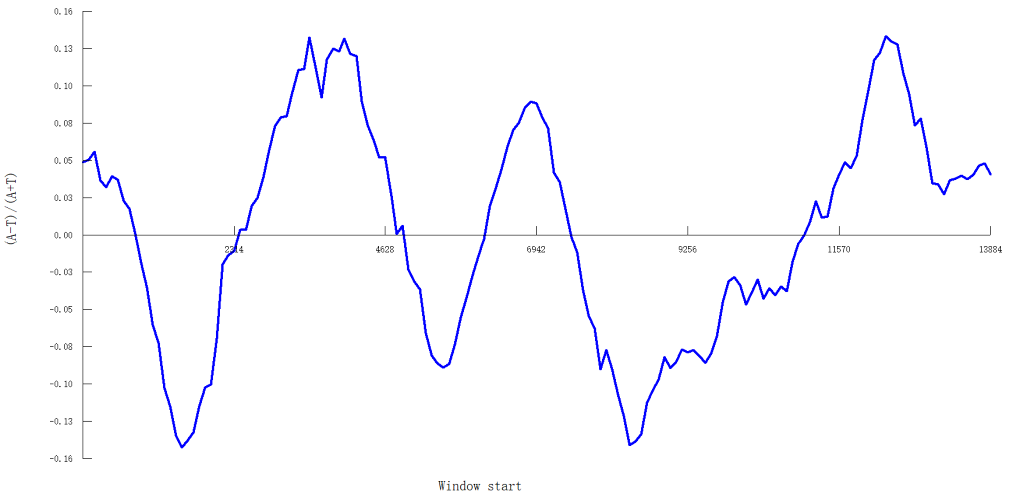


Figure S17. *Sinergasilus polycolpus* AT (manually to 1500 bp)

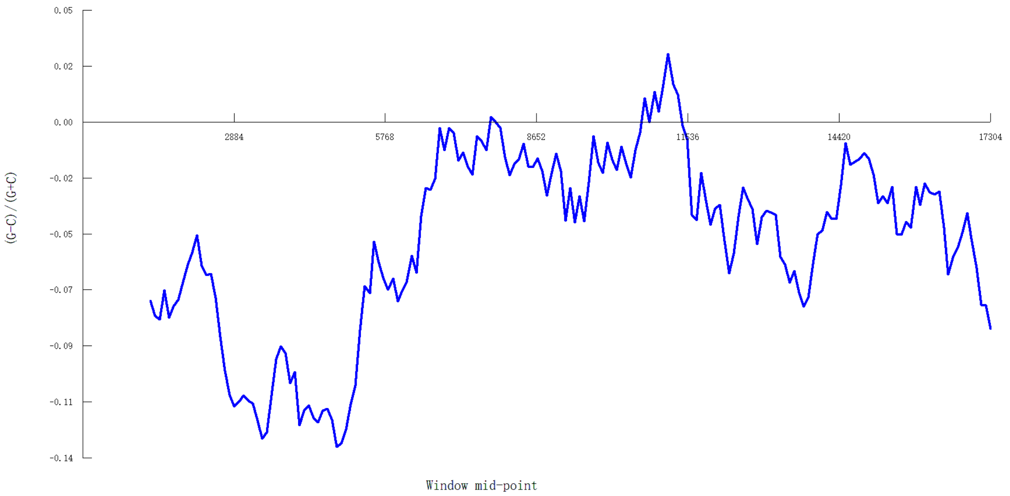


Figure S18. *Paracyclopina nana* GC

### Isopoda

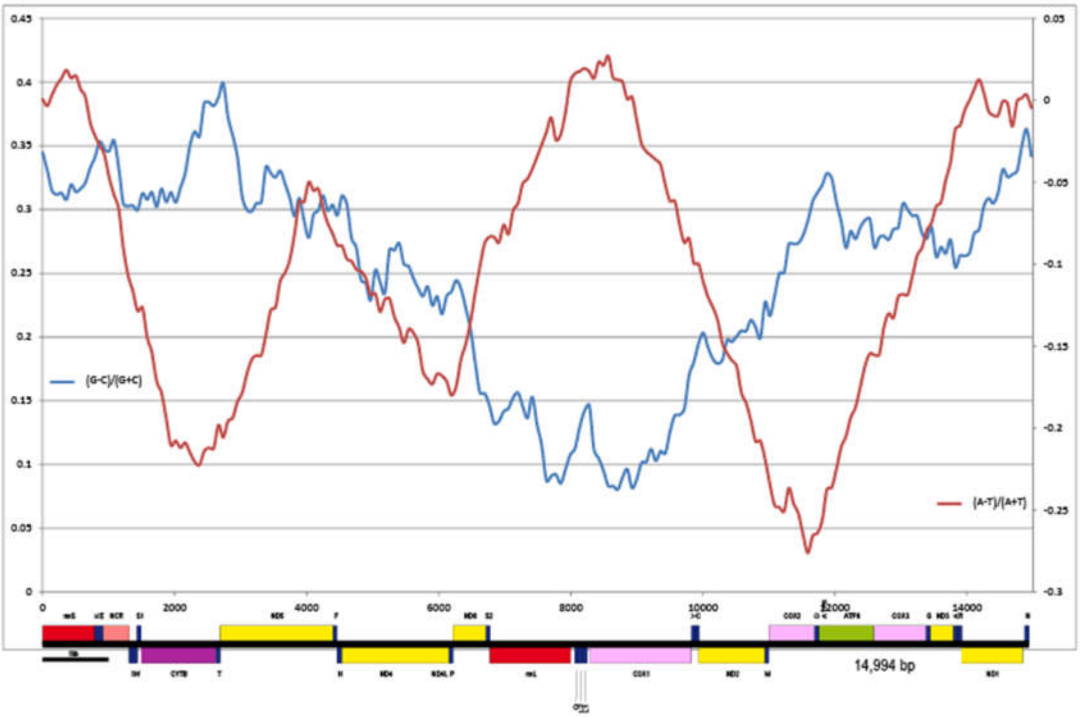


Figure S19. *Eophreatoicus sp.* Global GC Skew = 0.25049, Global AT Skew = -0.10440, Window size = 2202; Step size = 72.

### Amphipoda

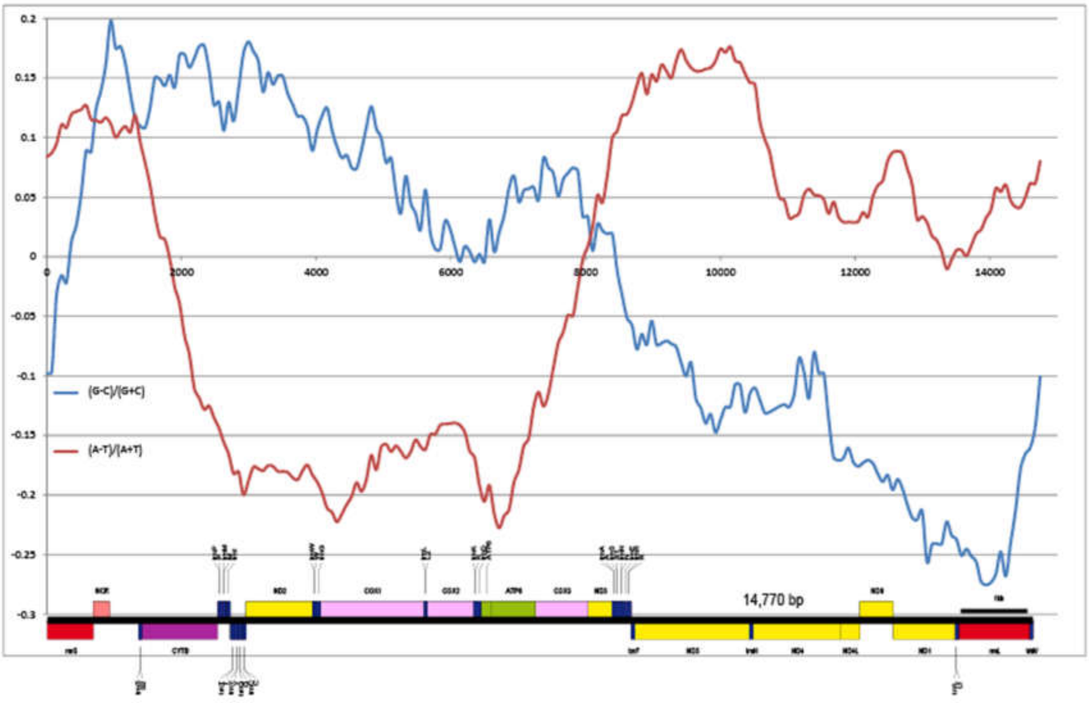

Figure S20. *Metacrangonyx ilvanus*, Global GC Skew = -0.01198, Global AT Skew = -0.01444, Window size = 1664; stepsize = 73.

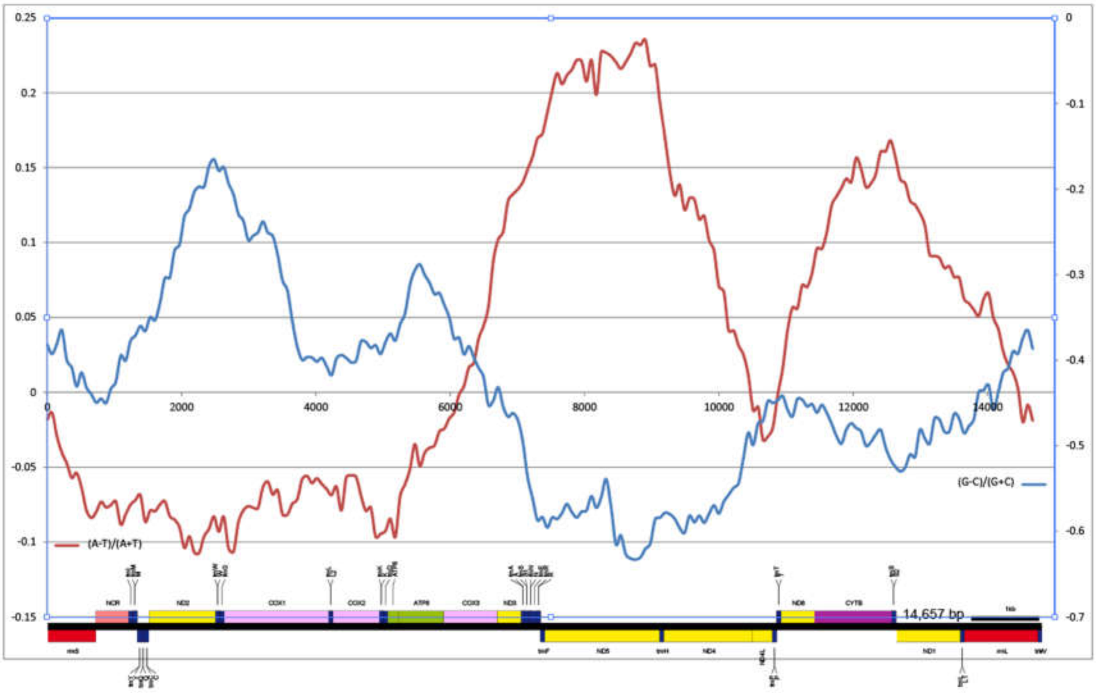


Figure S21. *Bahadzia jaraguensis*, Global GC Skew = -0.43082, Global AT Skew = 0.03682, Window size = 1756; stepsize = 73.

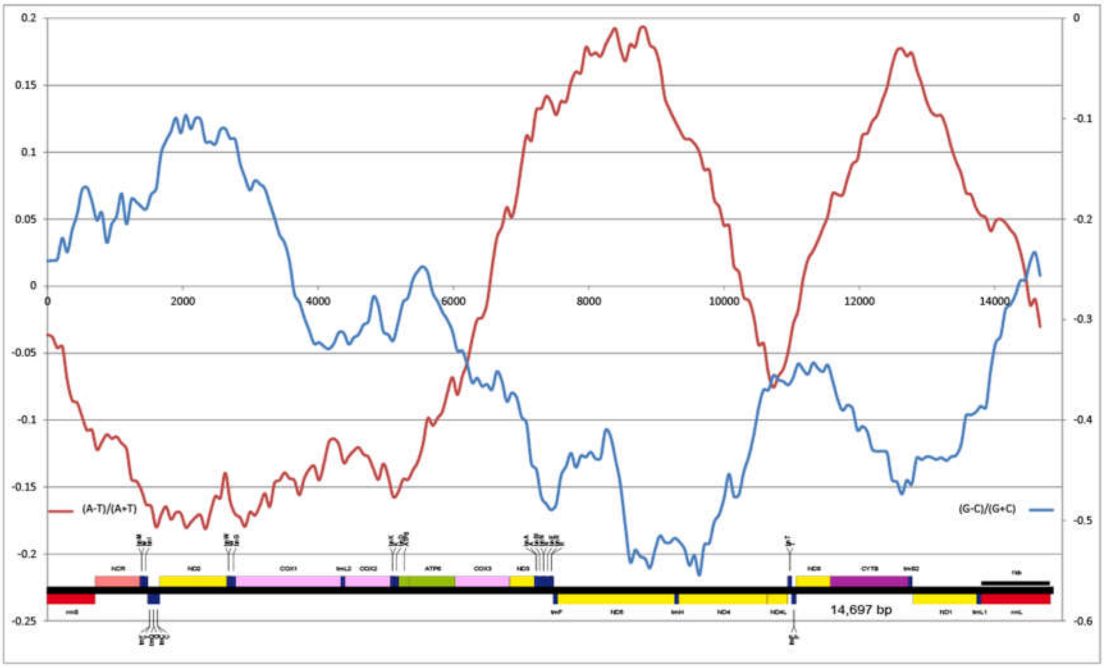


Figure S22. *Pseudoniphargus stocki* , Global GC Skew = -0.33547, Global AT Skew = -0.00649, Window size = 1910; stepsize = 73

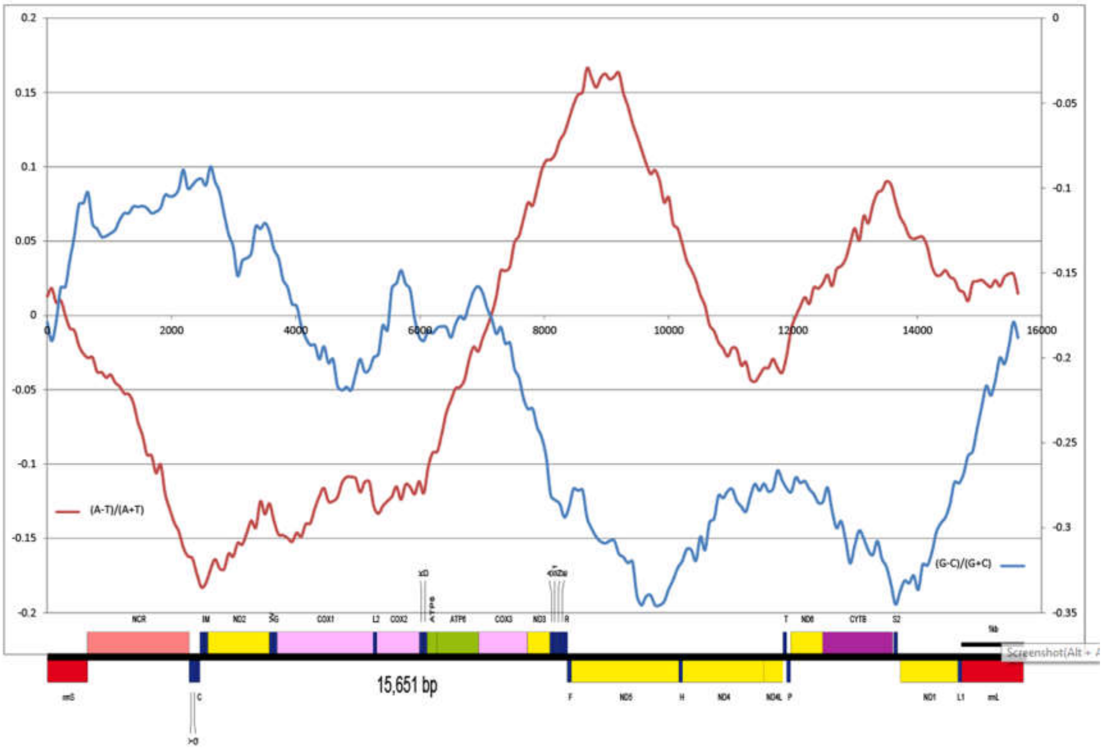


Figure S23. *Gammarus duebeni*, Global GC Skew = -0.22274, Global AT Skew = -0.01587, Window size = 2655; stepsize = 73.

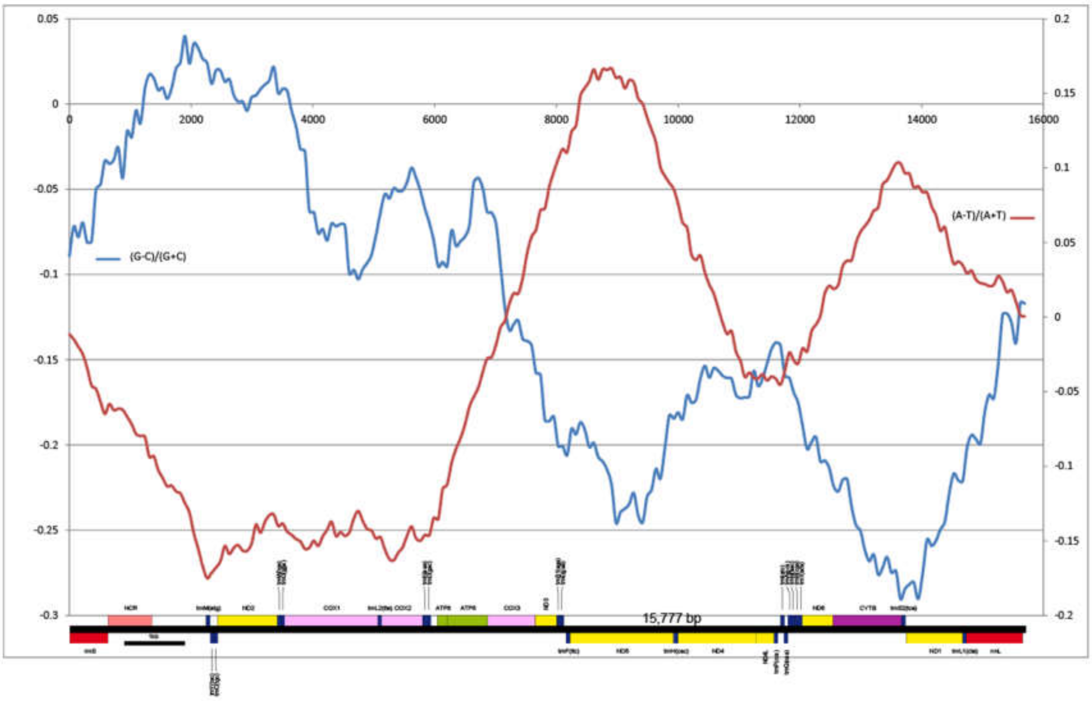


Figure S24. *Paryhale hawaiensis*, Global GC Skew = -0.1213, Global AT Skew = -0.021, Window size = 2702; stepsize = 73.

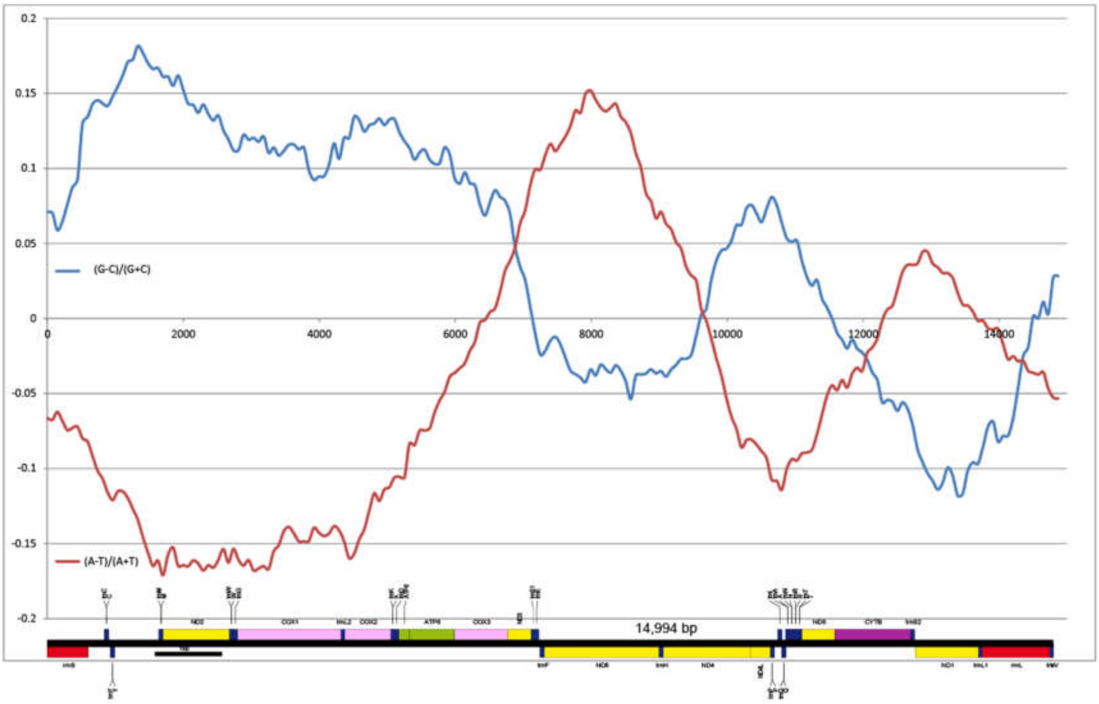


Figure S25. *Hyalella lucifugax*, Global GC Skew = 0.047, Global AT Skew = -0.042, Window size = 2702; stepsize = 74

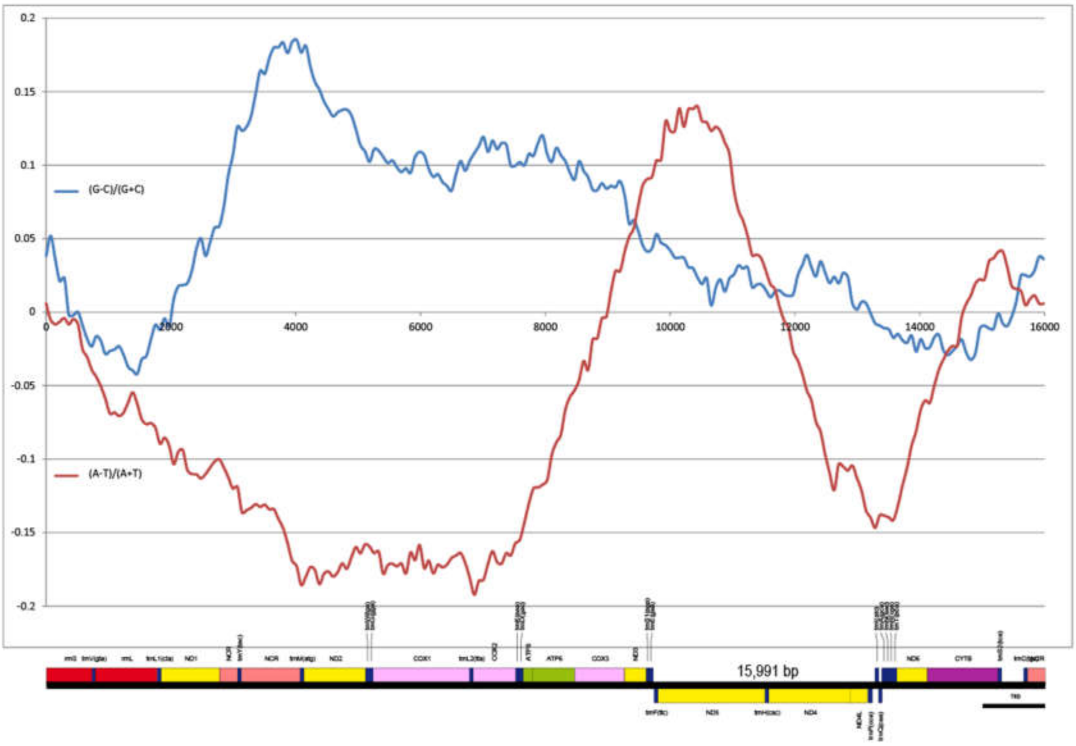


Figure S26. *Hyalella azteca*, Global GC Skew = 0.05194, Global AT Skew = -0.06570, Window size = 2702; stepsize = 73

### Decapoda

### Axiidea

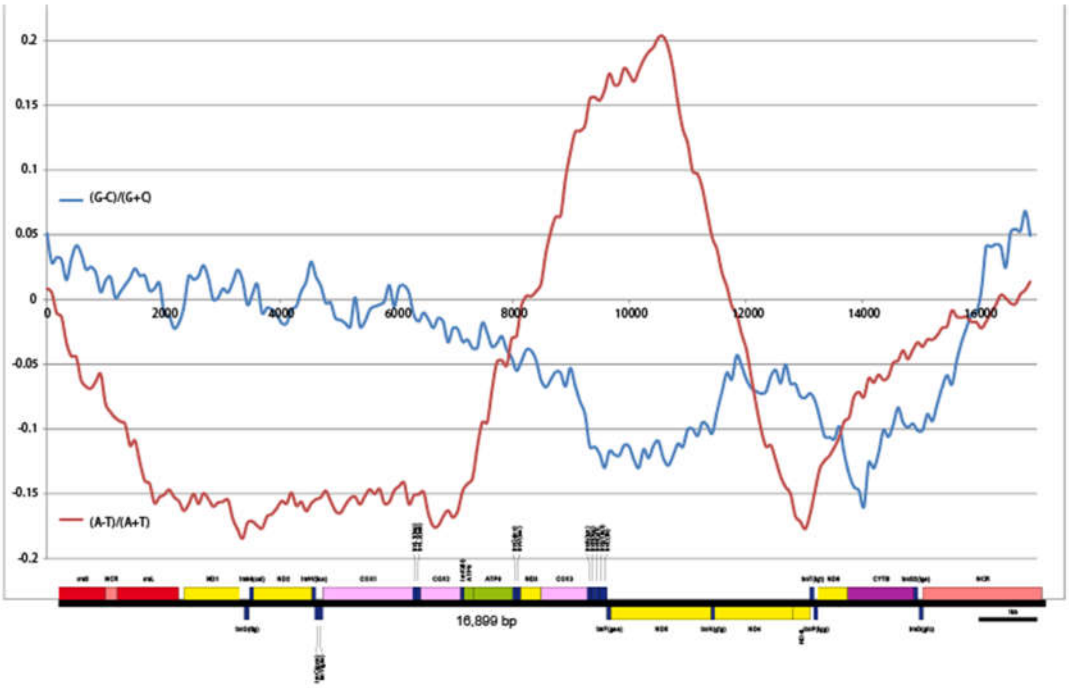


Figure S27. *Callianassa ceramica*, Global GC Skew = -0.03925, Global AT Skew = -0.05553

Window size = 2365; stepsize = 84.

### Stomatopoda

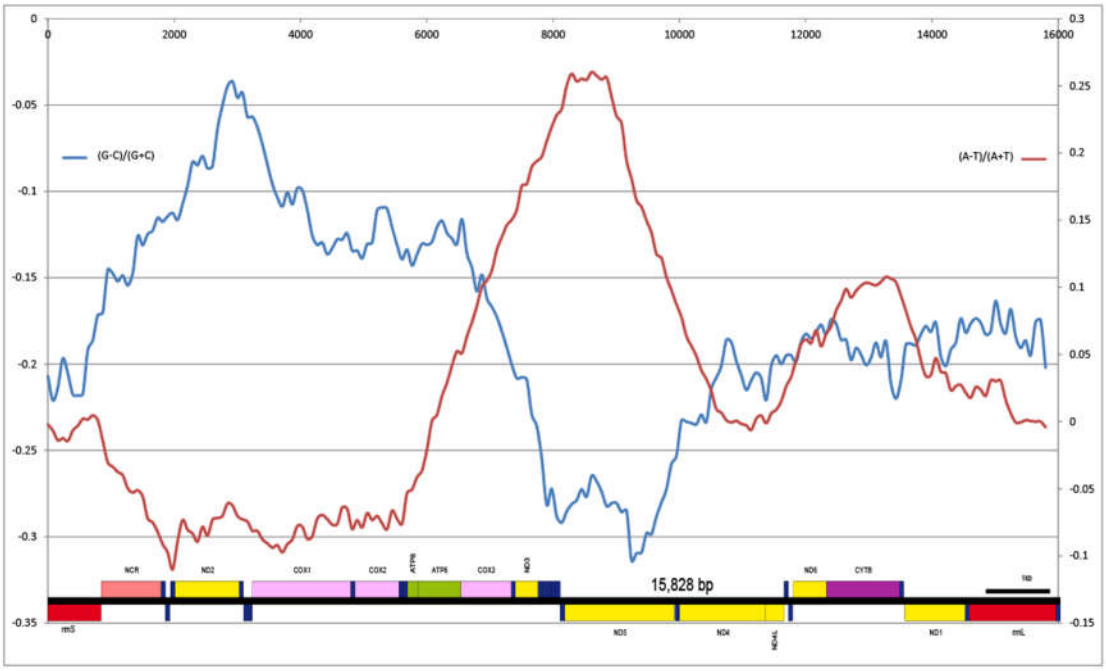


Figure S28. Cumulative skew plot for *Squilla empusa*.
